# Genome-wide screens identify calcium signaling as a key regulator of IgE^+^ plasma cell differentiation and survival

**DOI:** 10.1101/2021.03.02.433398

**Authors:** Rebecca Newman, Pavel Tolar

## Abstract

IgE antibodies protect against toxins and parasites, however, they also mediate allergic reactions. In contrast to other antibody isotypes, B cells switched to IgE respond transiently and do not give rise to long-lived plasma cells (PCs) or memory B cells. Although the intrinsic differences of IgE^+^ B cells have been linked to signaling by the IgE-B cell receptor (BCR), the molecular pathways controlling their behavior remain poorly understood. Here we employ whole-genome CRISPR screening to identify genes regulating IgE^+^ B cell proliferation, survival and differentiation into PCs. We show that IgE^+^ B cells are selectively suppressed by the IgE-BCR signaling to intracellular calcium, which inhibits PC differentiation and limits their lifespan after differentiation. Consequently, manipulation of calcium signaling *in vivo* enhances IgE^+^ PC responses. Insights from this pathway shed new light on the self-limiting character of IgE responses and open new avenues to eliminate IgE^+^ PCs in allergy.

## Introduction

B cells play crucial roles in the immune response by recognizing foreign antigens and eliciting appropriate host protective responses. Immunoglobulin class-switching to IgG, IgE and IgA is a major mechanism to diversify B cell responses and match antibody effector functions to immune challenge. Immunoglobulin E (IgE) antibodies, the least abundant of all antibody classes, have been thought to provide critical immunity to helminths (Fitzsimmons et al., 2014), but also potently protects from venoms (Marichal et al., 2013; Starkl et al., 2016) and cancers (Crawford et al., 2018). However, in modern humans, IgE is also frequently produced in response to innocuous environmental substances such as pollen, dust or common foods, underlying a spectrum of allergic diseases including life-threatening anaphylaxis, the prevalence of which continues to increase worldwide (Pawankar, 2014). As such, IgE production needs to be tightly regulated (Laffleur et al., 2015).

IgE responses, similar to other antibody isotypes, are initiated by antigen-mediated activation of B cells and their collaboration with cognate helper T cells. Helper T cells of the Th2-type promote class-switching of the activated B cells to IgE via the cytokine IL-4, which triggers DNA recombination at the immunoglobulin heavy chain locus. However, secretion of IgE antibodies by class switched B cells first requires expression and signaling activity of the membrane form of IgE as part of the cell-surface B cell receptor (in complex with the CD79A and CD79B subunits), indicating a role for IgE-BCR signaling at the plasma membrane(Achatz et al., 1997; Schmitt et al., 2020). Recently, the IgE-BCR was found to differ in its signaling capacity from the IgM- and IgG1-BCRs in that it initiates signaling in the absence of antigen, limiting B cell proliferation and stimulating differentiation into IgE^+^ PCs (Haniuda et al., 2016; Yang et al., 2012). This mirrors the low IgE^+^ B cell numbers, transient appearance, precocious differentiation into short lived PCs and the lack of memory cells *in vivo* (Erazo et al., 2007; He et al., 2013; Yang et al., 2012), arguing that signaling from the IgE-BCR critically regulates IgE production.

Despite the importance of IgE-BCR signaling, the precise differences compared to IgM- and IgG-BCR signaling and their impact on cellular outcomes remain to be fully understood. Although all membrane-bound immunoglobulin isotypes are associated with the transmembrane proteins CD79A and CD79B, which are required for BCR signal transduction, membrane IgG and IgE heavy chains also possess a cytoplasmic tail containing an evolutionarily conserved immunoglobulin tail tyrosine (ITT). The ITT is phosphorylated upon antigen engagement and has been implicated in facilitating elevated calcium and MAPK signaling through the signaling adaptor GRB2 (Engels et al., 2009). It is thought that IgE-BCRs have the additional ability to produce constitutive, low-level signaling in the absence of antigen binding, and this may occur in part through a unique extracellular region which enables binding to and signaling through the membrane adaptor CD19, as well as signaling through the cytoplasmic adaptor BLNK (Haniuda et al., 2016; Yang et al., 2016). All of these signaling pathways, however, are shared by antigen-stimulated IgG1^+^ cells, making it unclear how discrete fate decisions of IgE^+^ B cells are encoded. Proteins specifically binding to either the IgG1- (Liu et al., 2012) or IgE-BCR (Oberndorfer et al., 2006) have been described, but their physiological contributions to this conundrum remain unclear. Additional differences between IgG1^+^ and IgE^+^ cells and PCs have been described at the transcriptional level (Croote et al., 2018; Ramadani et al., 2019), but again how these arise and what their functional contribution is remains unknown and nor is the role of the paradoxically enhanced surface membrane IgE-BCR expression on IgE^+^ PCs, which contrasts with the decreased expression of membrane IgG1-BCR on IgG1^+^ PCs (He et al., 2013; Ramadani et al., 2017).

To better understand the mechanisms by which the IgE-BCR controls IgE responses, we employed whole-genome CRISPR screening and identified genes regulating the generation, proliferation, survival and PC differentiation of IgE^+^ B cells. These experiments confirmed the essential role of antigen-independent BCR signaling in IgE^+^ B cell biology. By comparing IgE^+^ cells to IgG1^+^ cells we identified common and IgE-specific pathways regulating cell death and PC differentiation. We show that IgE^+^ PC differentiation is driven by a class-common PI3K-mTOR signaling regulating levels of IRF4 protein. In contrast, IgE responses are selectively inhibited by calcium signaling, which inhibits PC differentiation and persists after IgE^+^ PC differentiation to culminate in their apoptosis. Targeting calcium signaling enhanced IgE^+^ PC numbers *in vivo*. Understanding IgE-selective pathways can provide insight into accumulation of IgE producing cells in allergic disease and offer new targets for treatments.

## Results

### Genome-wide CRISPR screens in primary B lymphocytes

To identify genes regulating responses of IgE^+^ B cells on a genome-wide scale, we performed pooled CRISPR screens using the mouse Brie CRISPR library comprising 78,637 sgRNAs targeting 19,674 genes, with 1000 non-targeting negative control sgRNAs (Doench et al., 2016). To enable screening with this library in primary mouse B cells without drug selection, we used Gibson assembly to clone the library into a lentiviral plasmid containing mCherry in place of a puromycin selection cassette (Supplementary Figure 1A) and confirmed that library complexity was well maintained in the Cherry Brie library (Supplementary Figure 1B, C). For each of our screens we used *ex vivo* isolated primary mouse B cells expressing Cas9-GFP from the Rosa26 locus (Platt et al., 2014). We infected the B cells with the lentiviral Cherry Brie library at a multiplicity of infection (MOI) of 0.3-0.5 and cultured the transduced B cells in the presence IL-4 on 40LB feeder cells (Nojima et al., 2011), which express CD40L and TNFSF13B (BAFF). Naïve follicular B cells subjected to this culture rapidly proliferate and express markers characteristic of GC B cells, referred to here as induced GC (iGC) cells, and undergo class-switching to IgG1 and IgE (Haniuda et al., 2016; Nojima et al., 2011). After six days in culture, we removed B cells from the feeders and cultured them without cytokines for two further days to promote plasma cell differentiation (Figure 1A). At the end of the culture, we flow sorted GFP^+^, mCherry^+^ IgE^+^ iGCs (IgE^+^, CD138^−^), IgG1^+^ iGCs (IgG1^+^, CD138^−^) and/or IgE^+^ PCs defined here as IgE^+^, B220^lo^, CD138^+^ cells. We then amplified and sequenced sgRNAs from each sorted population (sgRNA population) as well as from 293T cells used to produce the lentiviral library (sgRNA library). Library complexity was maintained across all three screens based on read count distribution for each sgRNA, with minimal guides having low or no read-counts (Supplementary Figure 1D-F). Using MAGeCK analysis (Li et al., 2014) to compare the abundance of sgRNAs in a given sorted population to the abundances in the original sgRNA library, we identified sgRNAs that were enriched or depleted in our populations of interest which are assigned positive or negative CRISPR scores respectively (Figure 1A).

**Figure 1:**
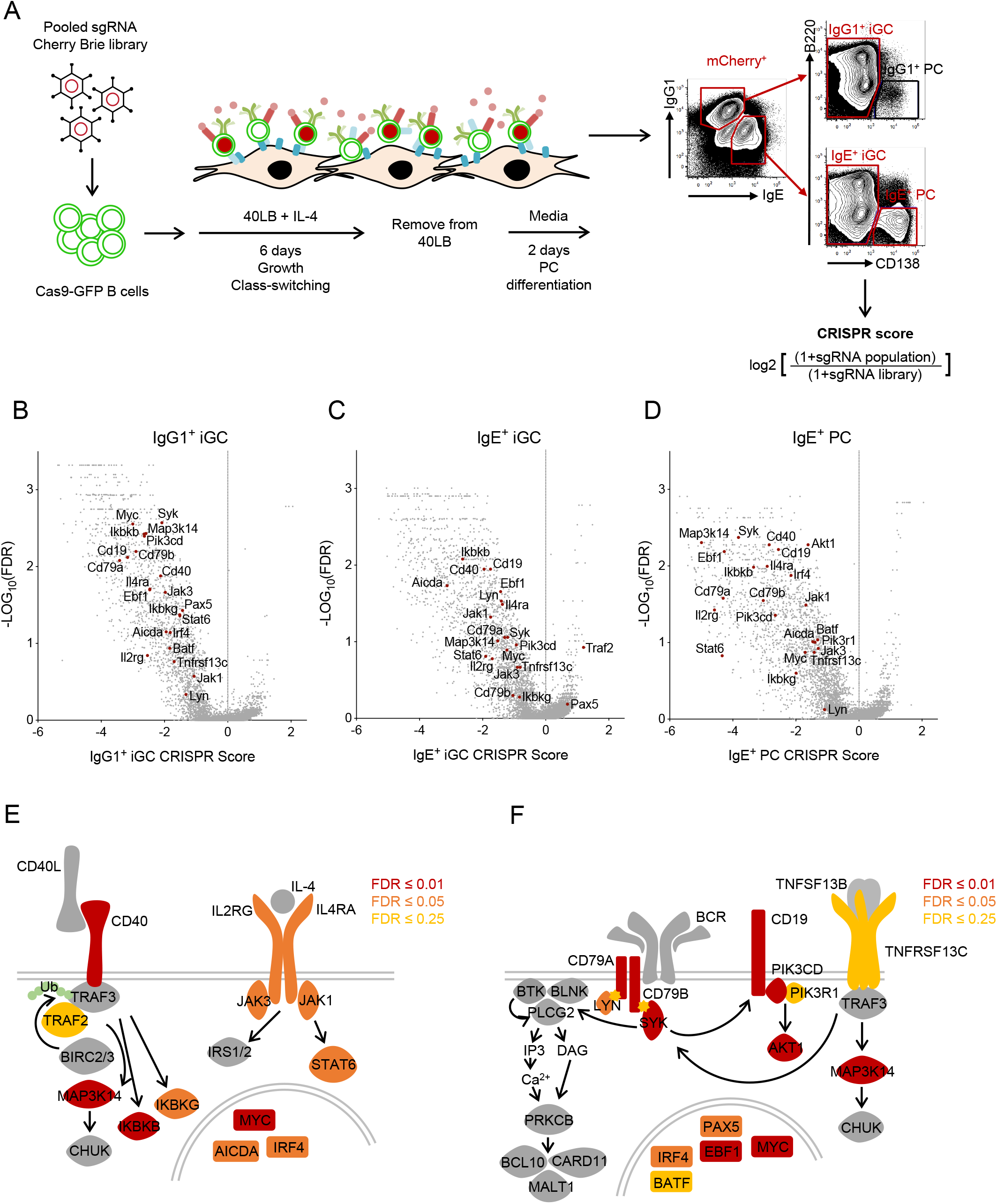
Genome-wide CRISPR screens in primary B lymphocytes. **A** Schematic of screen workflow. **B**, **C**, **D** Volcano plots of IgG1^+^ iGC (**B**), IgE^+^ iGC (**C**), IgE^+^ PC (**D**) CRISPR screens, showing CRISPR score (average sgRNA enrichment or depletion per gene) vs statistical significance corrected for false discover rate (FDR) for all genes (grey) and expected essential genes (dark red). **E**, **F** Schematics showing CD40 and IL-4 signaling pathways (**E**), BCR and TNFRSF13C signaling pathways (**F**), indicating minimum FDR (≤ 0.01 dark red, ≤ 0.05 orange, ≤ 0.25 yellow) across all screens for indicated gene products. For minimum FDR > 0.25, or where genes are not targeted in the CRISPR screen (PLCG2), signaling pathway components are indicated in grey.

To verify that our screening approach has been successful we compared the gene-averaged abundances of sgRNAs targeting 187 common essential genes (Wang et al., 2015), negative control non-targeting sgRNAs and all sgRNAs in the library. As expected, in all three populations, sgRNAs targeting common essential genes were heavily depleted (negative CRISPR scores), abundance of non-targeting control sgRNAs remained unchanged (CRISPR scores of close to zero), and the frequency of sgRNAs across the library showed a range of CRISPR scores (Supplementary Figure 1G-L).

RNA-sequencing (RNAseq) data confirmed that the cell-types generated using the *in vitro* B cell culture are transcriptionally distinct (Supplementary figure 2A, B) and express key transcripts which are associated with GC (*Bcl6*, *Pax5*) or PC (*Prdm1, Irf4, Xbp1*) fates (Supplementary figure 2C-G, Supplementary Table 1). The expression of Igh genes (Supplementary figure 2H-J) also validates our sorting strategy. Reflecting published data (Kräutler et al., 2017; Phan et al., 2006), IgG1^+^ PC are rare without the addition of antigen, and were therefore not included in this set of CRISPR screens. We found that stimulation of B cells after feeder removal using anti-Igκ F(ab’)_2_ as a surrogate antigen had no impact on IgE^+^ PC differentiation (Supplementary Figure 3A, B) but promoted IgG1^+^ PC differentiation in a dose dependent manner (Supplementary Figure 3C, D). We could recapitulate these findings using cultured SWHEL B cells (Phan et al., 2003) and soluble HEL antigen (data not shown).

To evaluate further the quality of our screens we inspected the top ranked essential genes for known signaling components downstream of the BCR, CD40, BAFF-R and IL-4R (Figure 1B-D). The results are summarized by schematics depicting the statistical significance of individual genes in these pathways in all three screens (Figure 1E, F). A shared requirement for these essential genes between the differentiated cells is also evident from comparative dot plots comparing CRISPR scores from IgE^+^ iGC and IgG1^+^ iGC screens (Supplementary Figure 4A) as well as comparison of IgE^+^ iGC and IgE^+^ PC screens (Supplementary Figure 4B).

### Novel regulators of B cell biology

In addition to identifying known regulators of BCR, CD40, BAFF-R and IL-4R driven B cell survival, proliferation and class-switching, we identified a number of genes previously not recognized as having roles in B cells. Among these was *Morc3*, a regulator of gene silencing by binding to H3K4me3 marked chromatin (Li et al., 2016), that is highly phosphorylated after BCR stimulation (Satpathy et al., 2015). We found it to be essential in both IgE^+^ and IgG1^+^ iGC cells in the screens (Supplementary Figure 5A), which was confirmed by using two sgRNAs distinct from the ones in the Cherry Brie library (Supplementary Figure 5B).

To test the role of *Morc3 in vivo*, we generated chimeras by adoptive transfer of GFP-Cas9-expressing hematopoietic stem cells (HSCs), which were lentivirally transduced with *Morc3* targeting sgRNA, into irradiated RAG1 KO mice (Supplementary Figure 5C). In these chimeras we identify *Morc3* KO cells using mCherry expression and found a substantial reduction in the proportion of mature KO B cells in the spleen compared to *Cd4* targeting control or to non-targeted mCherry^−^ cells (Supplementary Figure 5D-H). In contrast, targeting of *Morc3* had no effect on the proportions of CD4 T cells (Supplementary Figure 5I) indicating a previously unrecognized but critical role for *Morc3* specifically during B cell development.

### Unique regulators of IgE^+^ B cell biology

In order to identify non-redundant genes regulating IgE^+^ cells selectively, we compared the CRISPR scores for each gene between the IgE^+^ iGC screen and the IgG1^+^ iGC screens (Figure 2A, Supplementary Table 2). By selecting for genes that have a CRISPR score close to zero in the IgG1^+^ iGC screen, and thus a minimal role in these cells for survival, proliferation and/or differentiation, but have a CRISPR score ≤ −1.8, and FDR ≤ 0.25 in the IgE^+^ iGC screen we identified genes which essential only in the IgE^+^ iGC cells (Supplementary Table 3, key genes highlighted in blue Figure 2A). Similarly, selecting genes with a IgE^+^ iGC CRISPR score ≥ 0.95, and FDR ≤ 0.25, but have IgG1^+^ iGC CRISPR scores close to zero, we identified genes which negatively regulate IgE^+^ iGC cell, but not IgG1^+^ cells (Supplementary Table 3, key genes highlighted in red Figure 2A). We further removed genes that are not expressed in our populations of interest according to our RNAseq data (Supplementary Figure 2, Table 1). We also performed MAGeCK maximum-likelihood estimation (MLE) analysis (Li et al., 2015) to directly compare sgRNA frequencies between the IgE^+^ iGC and IgG1^+^ iGC populations. This allowed identification of selective regulators of IgE^+^ iGC cells based on statistical significance (FDR<0.25). Comparing the gene lists identified though CRISPR score comparison and MLE analysis indicates substantial overlap (Figure 2B, C). Functional enrichments in each gene list were identified using STRING analysis (Szklarczyk et al., 2019). This analysis revealed a key role for BCR signaling and cell death in regulating IgE iGCs (Figure 2B, C). Whilst apoptosis has previously been reported to limit IgE B cell expansion (Haniuda et al., 2016; Laffleur et al., 2015), importance of this pathway remains contentious (Yang et al., 2016) and will be explored further below.

**Figure 2:**
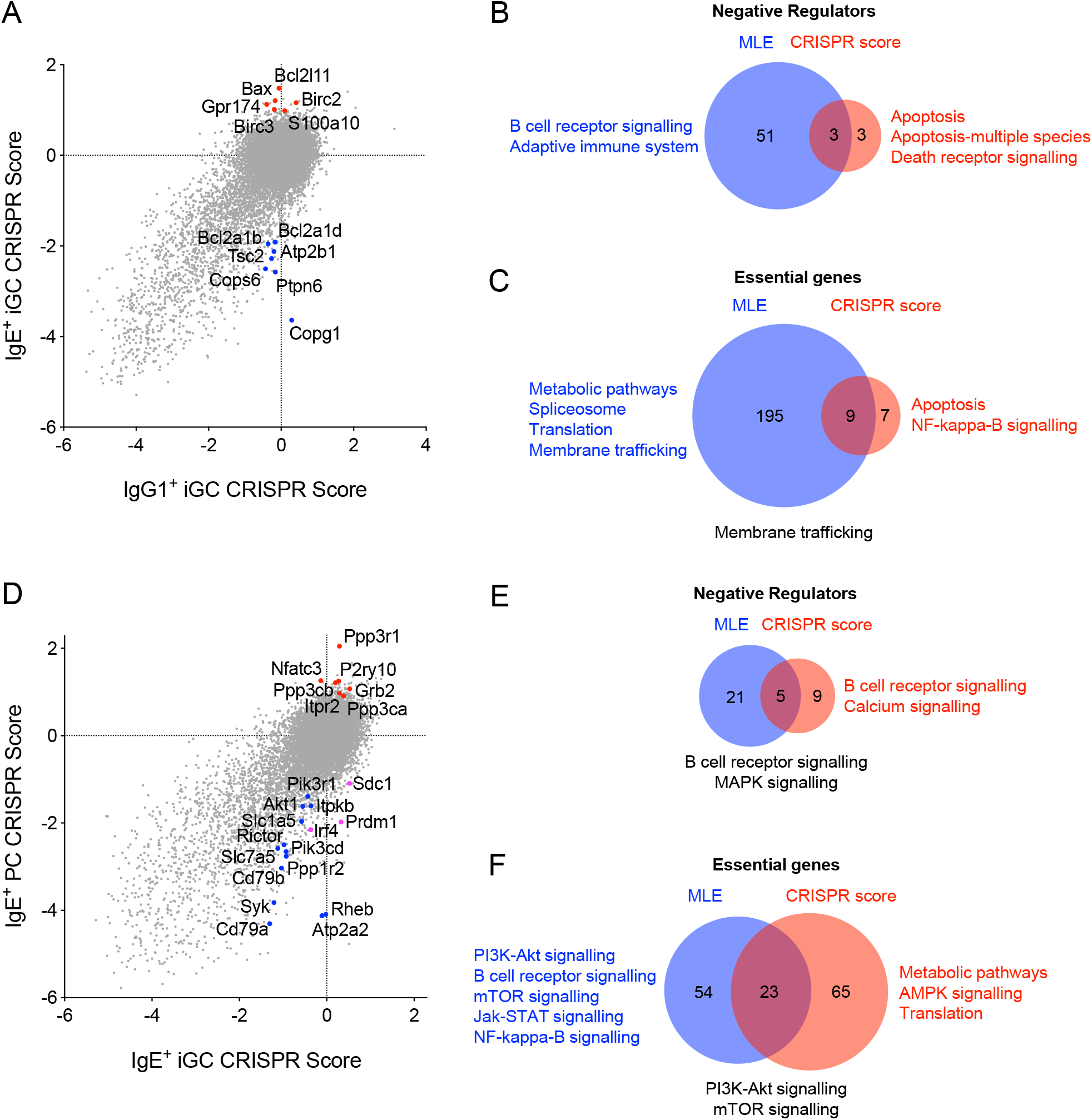
Identification of genes regulating IgE biology. **A** Comparative dot plot indicating key genes selectively essential in the IgE^+^ iGC screen. Each point represents a gene. Key IgE-specific essential genes (blue points) and IgE-specific negative regulators (red points) are indicated. **B**, **C** Venn diagrams indicating the proportional overlap between IgE^+^ iGC-specific gene lists generated using MLE analysis (FDR<0.25, blue circles) and CRISPR score cut-offs (red circles) for negative regulators (**B**) and essential genes (**C**). Functional enrichments in these gene lists were identified using STRING analysis (Szklarczyk et al., 2019). Key pathways are shown if applicable for MLE gene lists in blue, CRISPR score cut-offs in red or for overlapping gene lists in black. See Methods for CRISPR score and FPKM cut-offs defining these lists. **D** Comparative dot plot indicating genes essential in the IgE^+^ PC screen only. Each point represents a gene. Known regulators of PC differentiation are shown as pink points. Key IgE^+^ PC-specific essential genes (blue points) and IgE^+^ PC-specific negative regulators (red points) indicated. **E**, **F** Venn diagrams indicating the proportional overlap between IgE^+^ PC-specific gene lists generated using MLE analysis (FDR<0.25, blue circles) and CRISPR score cut-offs (red circles) for negative regulators (**E**) and essential genes (**F**). Functional enrichments in these gene lists identified using STRING analysis. Key pathways are shown if applicable as for MLE gene lists in blue, CRISPR score cut-offs in red or for overlapping gene lists in black.

To identify regulators of IgE^+^ PC differentiation and survival, we compared CRISPR gene scores from IgE^+^ iGCs with those of IgE^+^ PCs (Figure 2D-F). Using the same analysis as above we captured both *Irf4* and *Prdm1*, the central regulators of PC differentiation, and *Sdc1* which encodes CD138 used to sort this population (Figure 2D, pink dots, Supplementary Table 4). We validated that both *Irf4* and *Prdm1* were essential for IgE^+^ PCs independently of the screen (Supplementary Figure 6A and B respectively). STRING analysis identified PI3K-mTOR signaling amongst the top pathways essential for IgE^+^ PCs and calcium signaling amongst the top inhibitory pathways (Figure 2D, E, and Supplementary Table 4).

### The PI3K-mTOR pathway promotes IgE^+^ PC differentiation by enhancing IRF4 protein levels

PI3K signaling, stimulated by the IgE-BCR, has been proposed to promote IgE^+^ PC differentiation (Haniuda et al., 2016; Yang et al., 2016), but the mechanisms remains uncharacterized. Our data confirm the importance of this pathway and suggest that it regulates PC differentiation through mTOR. We first validated the role of mTOR pathways components using two further sgRNAs not in the sgRNA library in iGC cultures. In cells cultured in the same well we could distinguish and follow the differentiation, proliferation and survival of transduced KO (mCherry^+^) and non-transduced (mCherry^−^) WT cells. Both *Rheb*, a positive regulator of mTOR activity, and *Slc7a5*, an amino acid transporter required for full mTOR activation, were essential for IgE^+^ PC differentiation (Figure 2D, 3A, B). These requirements were shared with PC differentiation in IgG1^+^ cells induced by anti-Igκ F(ab’)_2_ acting as a surrogate antigen (Figure 3A, B). However, we also found *Slc7a5* was essential for IgE^+^ iGCs, but not for IgG1^+^ iGCs (Figure 3B). In addition, *Tsc2*, a negative regulator of *Rheb,* was also essential in IgE^+^ iGC cells, and showed a trend towards negatively regulating IgG1^+^ PC (Figure 2D, 3C). To confirm the importance of this, we cultured B cells in the presence of mTOR inhibitor, Rapamycin, and observed a specific loss IgE^+^ iGC cells as well as decreased IgE^+^ and IgG1^+^ PC differentiation (Figure 3D). We observed a specific loss of PCs when cells were cultured in the presence of Rapamycin for shorter time after class-switching had occurred (Supplementary Figure 7A). Similar results were observed when cells were cultured in the presence of Torin1 (data not shown). Thus, regulation of mTOR is selectively essential for IgE^+^ B cells while mTOR activity promotes both IgE^+^ and IgG1^+^ PC differentiation.

**Figure 3:**
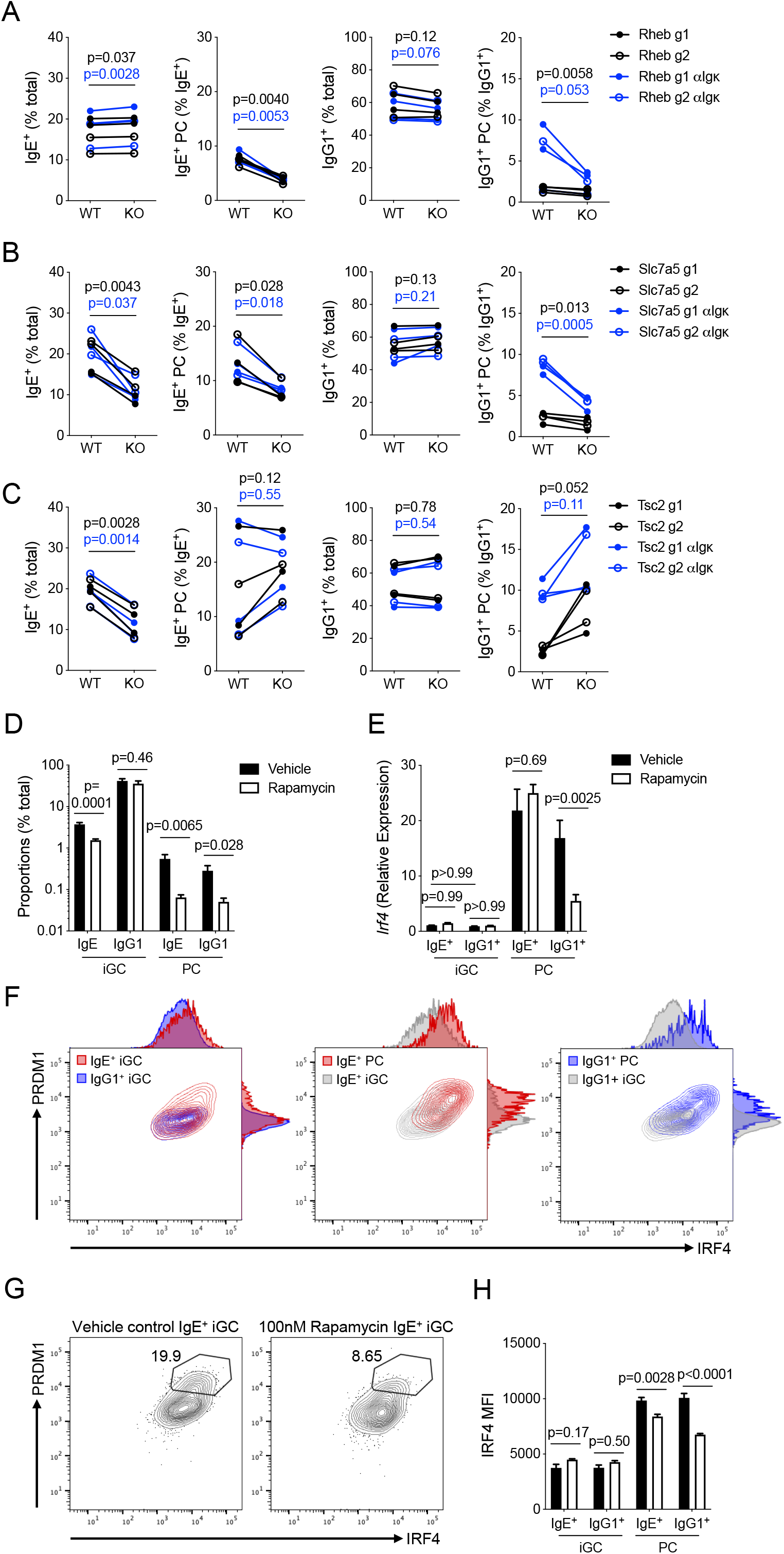
mTOR signaling drives IgE^+^ PC differentiation through post-transcriptional regulation of IRF4. **A** Screen validation data showing CRISPR KO of *Rheb* using 2 different sgRNAs (shown as open and closed circles). Conditions where 500ng/ml anti-Igκ F(ab’)_2_ was added for the final 2 days of culture are shown in blue. Proportions of IgE^+^ cells, IgE^+^ PCs, IgG1^+^ cells, and IgG1^+^ PCs (graphs from left to right) shown for WT (GFP^+^ mCherry^−^) and KO (GFP^+^ mCherry^+^) cells within the same culture well (connected by line). Data are pooled from 2 separate experiments. **B** Screen validation data showing KO of *Slc7a5* using 2 different sgRNAs, data are pooled from 2 separate experiments. (Plots as described for **A**). **C** Screen validation data showing KO of *Tsc2* using 2 different sgRNAs, data are pooled from 2 separate experiments. (Plots as described for **A**). Data in **A**-**C** were analyzed using paired t-tests. Significance values are indicated for cultures with (blue) and without (black) addition of anti-Igκ F(ab’)_2_. **D** Proportions of iGCs and PCs (as a % of live lymphocytes) with 100nM Rapamycin (white bars) or vehicle control (black bars) added for the final 5 days of B cell culture. N=8 biological replicates (data are pooled from 4 independent experiments) bars show mean ± SEM. Statistical significance (unpaired t-test) is indicated. **E** *Irf4* transcript abundance relative to *Hprt* and *Tbp* in sorted IgE^+^ and IgG1^+^ iGCs and PCs. Cells were cultured with 100nM Rapamycin (white bars) or vehicle control (black bars) for the final 5 days of B cell culture. Each bar represents 3 biological replicates, bars show mean ± SEM. Statistical significance (Two-way ANOVA with Sidak’s multiple comparisons test) indicated. (**F**) Representative flow cytometry contour plots indicating PRDM1 vs IRF4 protein expression in IgE^+^ iGC (IgE^+^, CD138^−^) shown in red, overlaid with IgG1^+^ iGC (IgG1^+^, CD138^−^) shown in blue (left plot), IgE^+^ iGC (grey), overlaid with IgE^+^ PC (IgE^+^, B220^lo^, CD138^+^) shown in red (middle plot), and IgG1^+^ iGC (grey) overlaid with IgG1^+^ PC (IgG1^+^, B220^lo^, CD138^+^) shown in blue (left plot), with adjunct histograms. Data are representative of 14 technical replicates, from three experiments. (**G**) Representative flow cytometry contour plots indicating PRDM1 vs IRF4 protein expression in IgE^+^ iGC cells. Cells were cultured with 100nM Rapamycin (right plot) or vehicle control (left plot) for the final 5 days of B cell culture. Gates indicate IRF4^hi^ PRDM1^hi^ population, numbers indicate proportion of cells within this gate. Data are representative of 3 biological replicates from 2 independent experiments. **H** Bar graph showing MFI of IRF4 in flow cytometry in IgE^+^ and IgG1^+^ iGC and PC. Cells were cultured with 100nM Rapamycin (white bars) or vehicle control (black bars) for the final 5 days of B cell culture. Each bar represents 3 biological replicates, bars show mean ± SEM. Statistical significance (Two-way ANOVA with Sidak’s multiple comparisons test) indicated. Data are representative of 3 independent experiments.

Mechanistically, the PI3K-mTOR pathway has been proposed to act upstream of IRF4, a key transcription factor needed to initiate PC differentiation by upregulating PRDM1 (Haniuda et al., 2016; Sciammas et al., 2006). However, the molecular mechanism by which mTOR regulates IRF4 remains unknown. mTOR activity was elevated in both IgE^+^ PCs and IgG1^+^ PCs in the culture, compared to total IgE^+^ and IgG1^+^ cells respectively, using staining for phosphorylated mTOR or its substrate S6 kinase (Supplementary Figure 7B, C). We found that the abundance of *Irf4* mRNA was increased in IgE^+^ and IgG1^+^ PCs but it was not different between IgE^+^ and IgG1^+^ iGC cells in our RNAseq data (Supplementary Figure 7D) and in RT-PCR quantification (Figure 3E). However, IRF4 protein was elevated in a subset of IgE^+^ iGC cells relative to IgG1^+^ iGC cells, in addition to the expected upregulation in IgE^+^ and IgG1^+^ PCs (Figure 3F). Culturing cells in the presence of Rapamycin resulted in loss of the IRF4-high PRDM1-high population of IgE^+^ iGC population and decreased IRF4 protein levels in both IgE^+^ and IgG1^+^ PCs (Figure 3G, H). However, whilst Rapamycin decreased *Irf4* transcript abundance in IgG1^+^ PC, it did not affect *Irf4* mRNA abundance in IgE^+^ iGCs or IgE^+^ PC compared to vehicle control (Figure 3E). Thus, mTOR signaling uniquely regulates protein levels of IRF4 post-transcriptionally in a fraction of IgE^+^ B cells, driving their PC differentiation.

### Calcium signaling negatively regulates IgE**^+^** PC differentiation

The top-ranked negative regulators of IgE^+^ PC differentiation were genes involved in BCR calcium signaling (Figure 2D, E, and Supplementary Table 4). These include *Grb2* which potentiates calcium signaling downstream of class-switched BCRs, the endoplasmic reticulum (ER)-localized calcium channel *Itpr2*, and members of the Calcineurin-NFAT pathway (*Ppp3r1*, *Ppp3cb*, *Ppp3ca*, *Nfatc3*). In contrast, *Itpkb*, which negatively regulates calcium signaling by converting IP_3_ to IP_4_, promoted IgE^+^ PC differentiation. We also observed essential roles in IgE^+^ iGC and PCs for genes controlling calcium homeostasis by removing calcium from the cytosol, such as the SERCA pump *Atp2a2* and the plasma membrane calcium pump *Atp2b1* (Figure 2A, D, Supplementary Table 3, 4). This finding was unexpected as calcium signaling has long been thought to promote PC differentiation (Engels and Wienands, 2018). We validated the importance of *Grb2* (Supplementary Figure 8A-C), *Itpr2* and the calcineurin-NFAT pathway by testing 2 new sgRNAs targeting each gene in individual assays. IP_3_R-2, was inhibitory for IgE^+^ PCs and IgG1^+^ PCs, while IP_3_R-3 (encoded by *Itpr3*) had only a small effect in IgG1^+^ PCs (Supplementary Figure 8D, E), indicating non-redundant roles for IP_3_Rs in B cells. *Ppp3r1* KO and *Nfatc3* KO resulted in a specific expansion of both IgE^+^ and IgG1^+^ PCs, with PPP3R1 having a stronger effect on IgE^+^ PC than on IgG1^+^ PCs (p=0.0055 when comparing fold change increase using paired t-test) (Figure 4A, B). Further supporting the importance of calcineurin in negatively regulating PC differentiation, deletion of *Crtc2*, which is activated following dephosphorylation by calcineurin (Altarejos and Montminy, 2011) also resulted in specific expansion of IgE^+^ PCs and antigen-stimulated IgG1^+^ PCs *in vitro*, although the latter did not reach statistical significance (Supplementary Figure 8F). In contrast, *Plcg2* was essential for both IgE^+^ or IgG1^+^ PCs, illustrating that the inhibitory effect of this pathway is selective to the mechanisms downstream of IP_3_ (Supplementary Figure 8G).

**Figure 4:**
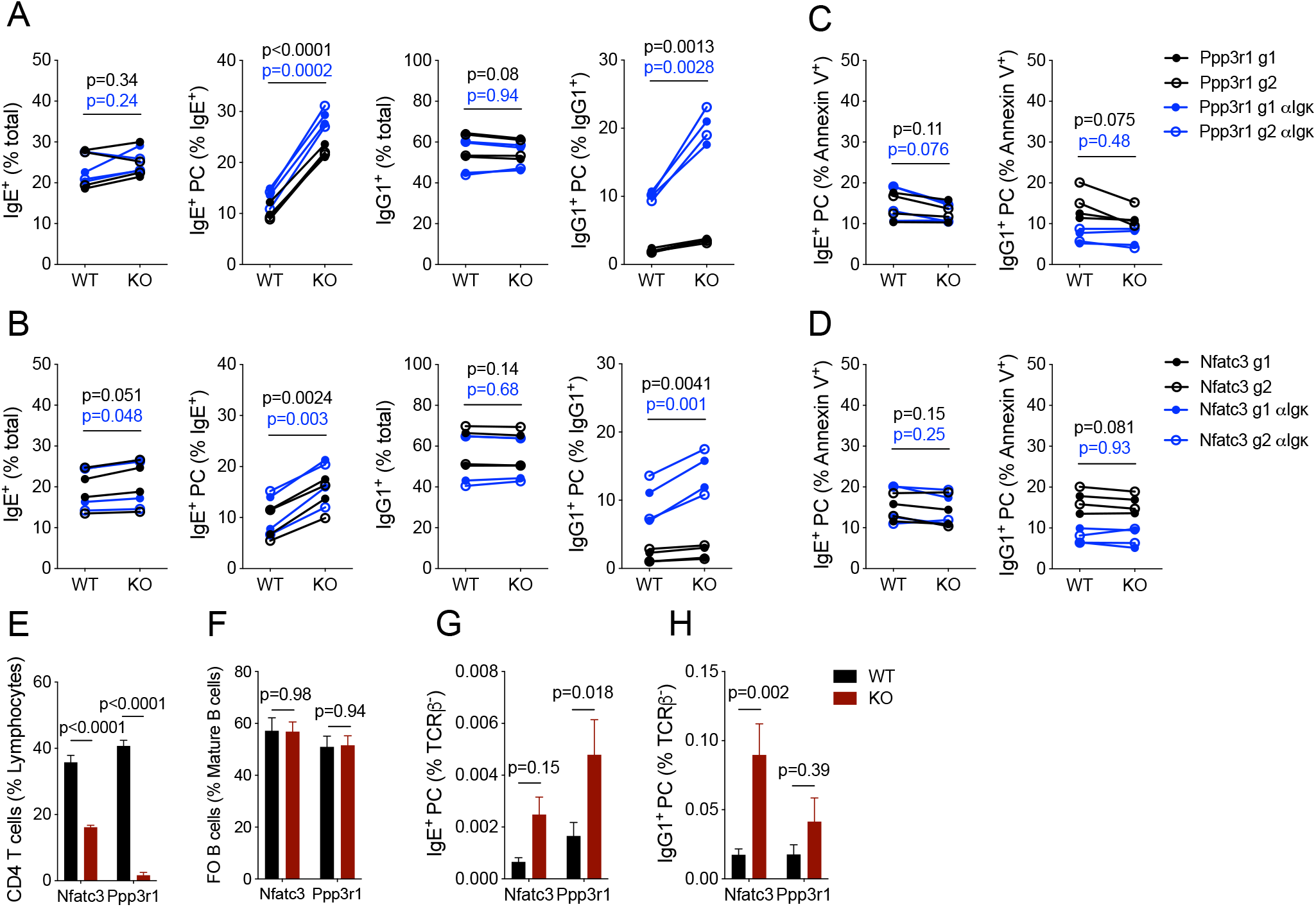
Negative regulation of PC differentiation by calcineurin signaling. **A** Screen validation data showing CRISPR KO of *Ppp3r1* using 2 different sgRNAs (shown as open and closed circles). Conditions where 500ng/ml anti-Igκ F(ab’)_2_ was added for the final 2 days of culture are shown in blue. Proportions of IgE^+^ cells, IgE^+^ PCs, IgG1^+^ cells and IgG1^+^ PCs (graphs from left to right) shown for WT (GFP^+^ mCherry^−^) and KO (GFP^+^ mCherry^+^) cells within the same culture well (connected by line). Data are pooled from 2 separate experiments. **B** Screen validation data showing CRISPR KO of *Nfatc3* using 2 different sgRNAs, are pooled from 2 separate experiments. (Plots as described for **A**). **C** Viability data showing KO of *Ppp3r1* using 2 different sgRNAs (shown as open and closed circles). Conditions where 500ng/ml anti-Igκ F(ab’)_2_ was added for the final 2 days of culture are shown in blue. Proportions of Annexin V^+^ IgE^+^ PCs, and IgG1^+^ PCs (graphs from left to right) shown for WT (GFP^+^ mCherry^−^) and KO (GFP^+^ mCherry^+^) cells within the same culture well (connected by line). Data are pooled from 2 separate experiments. **D** Viability data showing KO of *Nfatc3* using 2 different sgRNAs, data are pooled from 2 separate experiments. (Plots as described for **C**). Data in **A**-**D** were analyzed using paired t-tests. Significance values are indicated for cultures with (blue) and without (black) addition of anti-Igκ F(ab’)_2_. **E-H** Splenic CD4 T cell proportions (**E**), Splenic Follicular B cell proportions (**F**), draining LN IgE^+^ PC proportions (**G**) and draining LN IgE^+^ PC proportions (**H**) at day 7 in NP-CGG in Alum immunized CRISPR Cas9 chimeras comparing WT (GFP^+^ mCherry^−^) as black bars and KO (GFP^+^ mCherry^+^) as red bars. Data are pooled from 2 separate experiments, each bar is representative of 8 mice, bars represent mean ± SEM. Statistical significance (Two-way RM ANOVA with Sidak’s multiple comparisons test) is indicated.

To understand whether the calcineurin-NFAT pathway regulates PC differentiation or survival, we analyzed the proportion of Annexin V^+^ cells in *Ppp3r1* and *Nfatc3* CRISPR KO. The proportions of Annexin V^+^ PC were not different between either *Ppp3r1*- or *Nfatc3*-sufficient and deficient cells (Figure 4C, D). We conclude that the calcineurin-NFAT pathway primarily regulates PC differentiation rather than cell death.

To validate our findings in an *in vivo* setting we again generated lentiviral chimeras, as described above (Supplementary Figure 5C). Upon deletion of *Nfatc3* and *Ppp3r1* we observed a loss of CD4 T cells in the mCherry^+^ compartment as expected (Canté-Barrett et al., 2007; Neilson et al., 2004), but proportions of mature B cells were unchanged (Figure 4E, F). At day 7 following subcutaneous immunization of the chimeras using NP-CGG in Alum, we observed an increased proportion of GC B cells in the draining LNs (dLNs) in both the *Nfatc3* and *Ppp3r1* deficient mCherry^+^ compartment (Supplementary Figure 8H-J). There was an increase in the overall proportion of PCs amongst *Nfatc3* deficient cells (Supplementary Figure 8K) with a stronger increase in IgG1^+^ PCs (Figure 4G, H). In contrast, *Ppp3r1* KO resulted in a selective increase of IgE^+^ PCs (Figure 4G). As these chimeras have both WT and KO cells, these effects are B cell intrinsic. Thus, calcium signaling through calcineurin-NFATC3 restricts IgE^+^ and IgG1^+^ PC numbers both *in vitro* and *in vivo* and calcineurin itself is a stronger inhibitor of IgE^+^ cells than IgG1^+^ cells.

### Apoptosis is a key regulator of IgE**^+^** B cell and PC biology

Our screen data suggest that survival of IgE^+^ iGCs and IgE^+^ PCs is critically regulated by the balance of specific BCL2-family proteins, namely pro-apoptotic BCL2L11 (BIM) and BAX, and anti-apoptotic BCL2A1B and D (Figure 2, Supplementary Figure 4B and Supplementary Table 3). To verify the importance of this pathway we again tested targeting *Bcl2l11* and *Bcl2a1d* in individual assays. Deletion of *Bcl2l11* resulted in a specific expansion of IgE^+^ cells (Figure 5A) and IgE^+^ PCs (Figure 5B), whilst loss of *Bcl2a1d* resulted in loss of both IgE^+^ and IgG1^+^ cells (Figure 5C). To understand the role of *Bcl2l11* in more detail, we analyzed BCL2L11 protein levels and observed increased BCL2L11 staining by flow cytometry in IgE^+^ iGC cells when compared to IgG1^+^ iGC cells, with even higher levels in IgE^+^ PCs at day 8 of the culture (Figure 5D, E). To confirm these findings *in vivo*, we immunized mice subcutaneously with NP-CGG in Alum and found increased BCL2L11 expression in IgE^+^ GC B cells compared to IgG1^+^ GC B cells, and a similar trend in PCs (Figure 5F, G). Further supporting the role of apoptosis in limiting IgE^+^ B cells, we observed an increased proportion of Annexin V^+^ IgE^+^ cells at the end (day 8) of the culture indicating loss of viability relative to IgG1^+^ cells (Figure 5H, I). Furthermore, the proportion of active caspase 3^+^ cells, was higher in IgE^+^ iGC cells compared to IgG1^+^ iGC cells at day 6 and 8 of the culture and was further increased in IgE^+^ PCs at day 8 (Figure 5J, K). In addition, the proportion of active caspase 3^+^ cells was higher in IgE^+^ PCs when compared to IgG1^+^ PCs from mice immunized subcutaneously with NP-CGG in Alum, with a similar trend observed for GC B cells (Figure 5L, M). Altogether, these data suggest that compared to IgG1^+^ cells IgE^+^ cells undergo increased apoptosis both *in vitro* and *in vivo* and that this apoptosis involves BCL2L11 and culminates in PCs.

**Figure 5:**
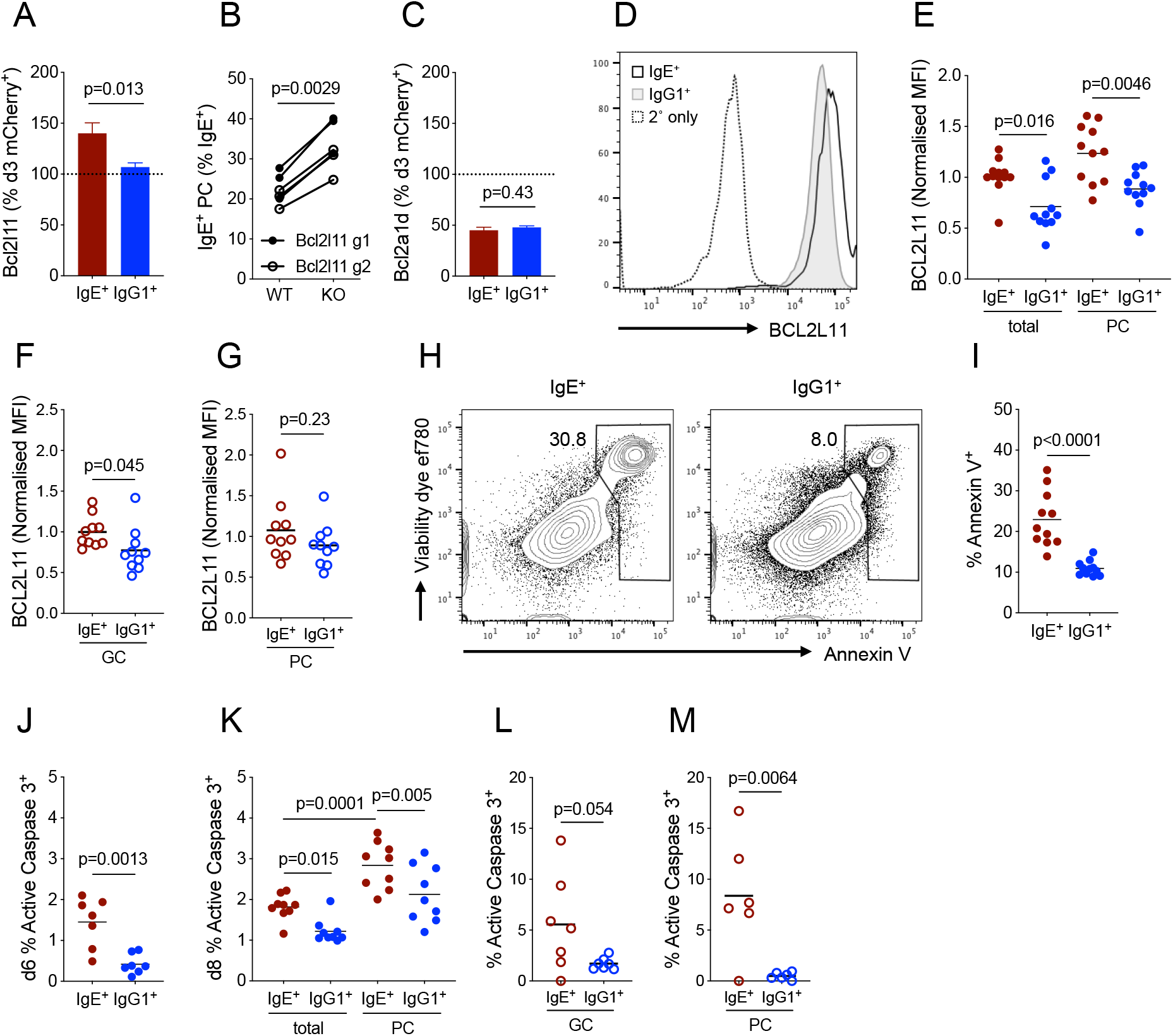
Identification of genes regulating IgE^+^ B cell apoptosis. **A** Validation of *Bcl2l11* CRISPR KO effect on IgE^+^ and IgG1^+^ cell numbers. Bars show mean ± SEM, statistical significance (unpaired t-test) is indicated. Validation shows data as mCherry^+^ IgE^+^ and IgG1^+^ cells at day 8 of B cell culture relative to proportions of mCherry^+^ B cells at day 3 of the culture. Data shown are pooled from KO effects of 2 sgRNAs per gene, from 2 experiments. **B** IgE^+^ PC proportions shown for WT (GFP^+^ mCherry^−^) and CRISPR KO of *Bcl2l11* (GFP^+^ mCherry^+^) cells within the same culture well (connected by line), using 2 different sgRNAs (shown as open and closed circles). Data are pooled from 2 separate experiments. **C** Validation of *Bcl2a1d* CRISPR KO, (plot as described for **A**). Data shown are pooled from KO effects of 2 sgRNAs per gene, from 1 experiment. **D** Representative FACS plot showing BCL2L11 expression in IgE^+^ (black line) and IgG1^+^ (Grey filled histogram) cells from day 8 of *in vitro* culture. Secondary antibody staining only is shown as a dashed line. **E** Normalized median fluorescence intensity for BCL2L11 staining using flow cytometry in IgE^+^ and IgG1^+^ cells and PC from day 8 of *in vitro* culture. Each dot represents a technical replicate, line indicates mean, data is pooled from 3 separate experiments. Statistical significance (One-way ANOVA with Tukey’s multiple comparisons test) indicated. **F, G** Normalized median fluorescence intensity for BCL2L11 staining using flow cytometry in IgE^+^ and IgG1^+^ GC B cells (**F**) and IgE^+^ and IgG1^+^ PCs (**G**) from mice immunized subcutaneously with NP-CGG in Alum (day 7 post-immunization). Each dot represents a biological replicate, line indicates mean. Data are pooled from 2 independent experiments; data were analyzed using an unpaired t-test. **H** Representative FACS plots indicating percentage of total IgE^+^ or IgG1^+^ cells (including iGCs and PCs) from day 8 of *in vitro* culture within Annexin V^+^ gate. Numbers on plots indicate the percentage of cells within the gate. **I** Proportion of Annexin V^+^ IgE^+^ and IgG1^+^ cells at day 8 of *in vitro* culture. Each dot represents a technical replicate, line indicates mean. **J** Proportion of active caspase-3^+^ IgE^+^ and IgG1^+^ cells from day 6 of *in vitro* culture. Each dot represents a technical replicate, line indicates mean. Data are pooled from 2 experiments. Statistical significance (unpaired t-test) shown. **K** Proportion of active caspase-3^+^ IgE^+^ and IgG1^+^ cells and PCs from day 8 of *in vitro* culture. Each dot represents a technical replicate, line indicates mean. Data are pooled from 2 experiments. Statistical significance (One-way ANOVA with multiple comparisons test) indicated. **L**, **M** Proportion of active caspase-3^+^ cells in IgE^+^ and IgG1^+^ GC cells (**L**) and IgE^+^ and IgG1^+^ PCs (**M**) from mice immunized subcutaneously with NP-CGG in Alum (day 7 post-immunization). Each dot represents a biological replicate, line indicates mean. Data are pooled from 2 independent experiments; data were analyzed using an unpaired t-test.

### Calcium downstream of the IgE-BCR results in loss of IgE^+^ B cells and PCs

To understand what triggers the apoptosis in IgE^+^ cells, we focused on the role of BCR signaling. We validated CRISPR screen data indicating that IgE^+^ iGC cells selectively required *Ptpn6* (that encodes for the inhibitory tyrosine phosphatase SHP1) (Figure 6A). This result was consistent with BCR kinase signaling providing pro-apoptotic signals in IgE^+^ B cells. Indeed, treating cells for the final 48 hours of culture with the Syk inhibitor (BAY 61-3606) resulted in increased proportions, and decreased cell death of IgE^+^ iGC cells, but decreased proportions of IgG1^+^ iGC cells with increased cell death (Figure 6B-E). Syk inhibition also resulted in decreased cytosolic calcium levels in both subsets (Figure 6F, G). More specifically, we implicated BCR signaling driving calcium release from the ER as important for apoptosis. Deletion of *Grb2* resulted in expansion of IgE^+^ B cells, but loss of *Grb2* was detrimental to antigen-dependent IgG1^+^ PC differentiation and/or survival (Supplementary Figure 8A), a phenotype that may result from GRB2 potentiation of MAPK signaling (Engels et al., 2009). However, IgE^+^ PCs were negatively regulated by GRB2 (Supplementary Figure 8A), possibly because MAPK signaling may be specific downstream of the antigen-bound IgG1 receptor, as it was not picked up in our IgE^+^ PC screens (Figure 2D and Supplementary Figure 4B). Consistently, analysis of immunized *Grb2*^fl/fl^*Cd79a*^cre/+^ mice indicated that Grb2 KO mice had increased proportions of IgE^+^ GC B cells (Supplementary Figure 8B) and decreased proportions of IgG1^+^ GC B cells (Supplementary Figure 8C) compared to and *Grb2*^fl/fl^ littermate controls. While *Plcg2* promoted apoptosis in IgE^+^ PCs (Supplementary Figure 9A), it was also required by both IgE^+^ and IgG1^+^ PC (Supplementary figure 8G), indicating that signaling downstream of PLCG2 plays a role both in PC differentiation, in-keeping with published data (Hikida et al., 2009), but also promotes apoptosis in IgE^+^ PCs. Finally, CRISPR KO of *Itpkb*, a negative regulator of calcium signaling resulted in a loss of IgE^+^ cells, and a decrease in IgE^+^ PCs, which again was associated with enhanced apoptosis in PCs (Supplementary figure 9B, C). Thus, mechanisms that enhance calcium signaling downstream of the IgE-BCR act to limit IgE^+^ B cell responses by promoting apoptosis in both IgE^+^ B cells and PCs. This BCR-calcium pro-apoptotic pathway shares the early signaling components with the BCR-calcium-calcineurin-NFAT pathway, which inhibits PC differentiation, but is genetically distinct in the downstream components.

**Figure 6:**
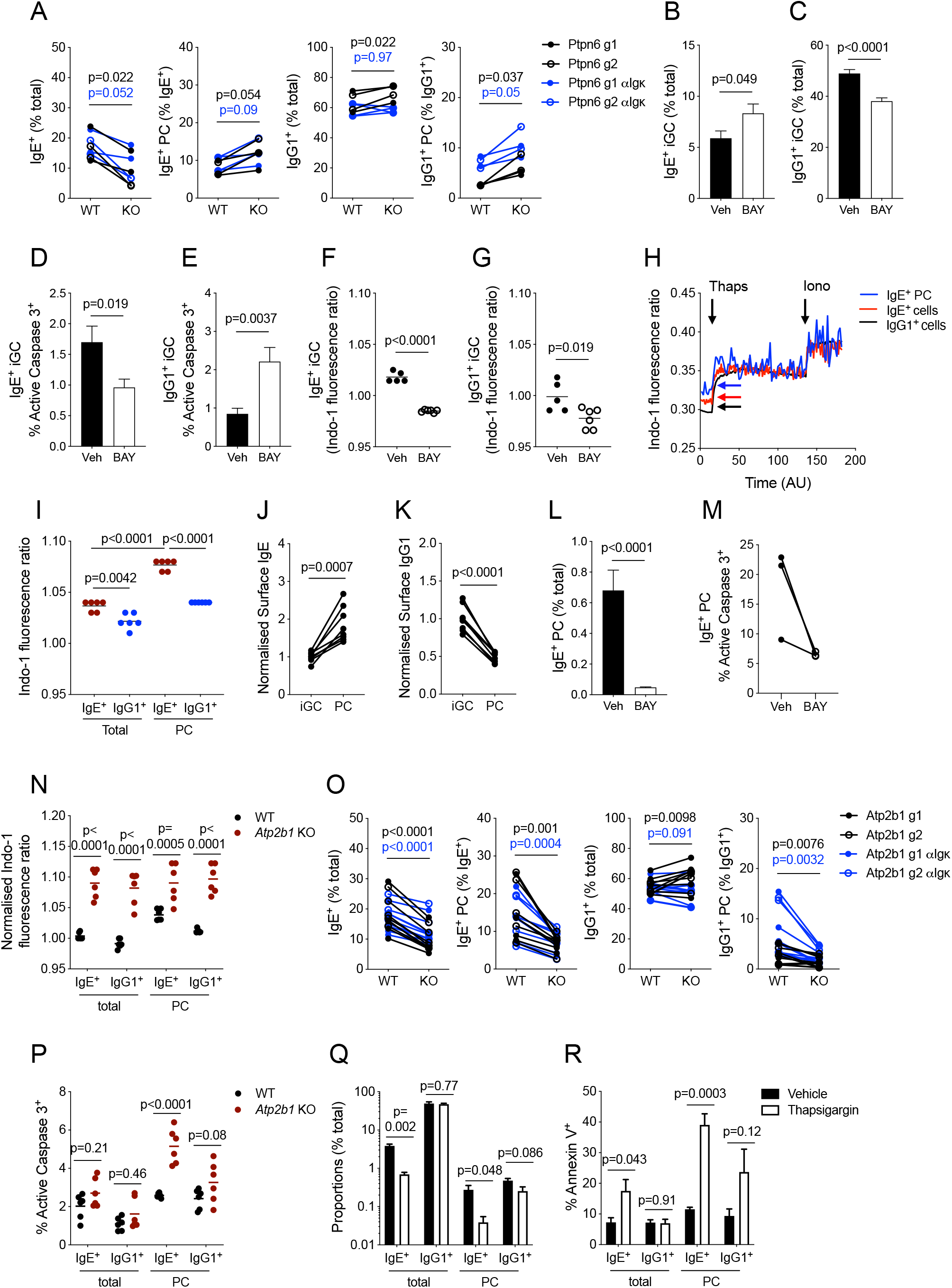
Calcium signaling regulates apoptosis of IgE^+^ cells. **A** Screen validation data showing KO of *Ptpn6* using 2 different sgRNAs (shown as open and closed circles). Conditions where 500ng/ml anti-Igκ F(ab’)_2_ was added for the final 2 days of culture are shown in blue. Proportions of IgE^+^ cells, IgE^+^ PCs, IgG1^+^ cells, and IgG1^+^ PCs (graphs from left to right) shown for WT (GFP^+^ mCherry^−^) and KO (GFP^+^ mCherry^+^) cells within the same culture well (connected by line), data are pooled from 2 separate experiments. Data in **A** were analyzed using paired t-tests. Significance values are indicated for cultures with (blue) and without (black) addition of anti-Igκ F(ab’)_2_. **B**, **C** Proportions of IgE^+^ iGCs (**B**) and IgG1^+^ iGCs (**C**) (as a % of live lymphocytes) treated with 1μM of the Syk inhibitor BAY 61-3606 (white bars) or vehicle control (black bars) added for the final 2 days of B cell culture (n=10-11, data pooled from 4 separate experiments) bars show mean ± SEM. Statistical significance (unpaired t-test) is indicated. **D**, **E** Proportions of active caspase3^+^ IgE^+^ iGCs (**D**) and IgG1^+^ iGCs (**E**) with 1μM of the Syk inhibitor BAY 61-3606 (white bars) or vehicle control (black bars) added for the final 2 days of B cell culture (n=10-11, data pooled from 4 separate experiments) bars show mean ± SEM. Statistical significance (unpaired t-test) is indicated. **F, G** Indo-1 fluorescence ratios in IgE^+^ iGCs (**F**) and IgG1^+^ iGCs (**G**) treated with 1μM of the Syk inhibitor BAY 61-3606 (open circles) or vehicle control (closed circles) added for the final 2 days of B cell culture (each point represents a technical replicate, data are pooled from 2 separate experiments) Statistical significance (unpaired t-test) is indicated. **H** Baseline calcium (indicated by arrows), followed by calcium flux induced by treatment of cells with 1uM thapsigargin (addition indicated by arrow labelled Thaps), followed by 5μg/ml Ionomycin (addition indicated by arrow labelled Iono) in IgG1^+^ cells (black line), IgE^+^ cells (red line) and IgE^+^ PCs (blue line). Calcium levels measured using flow cytometry as Indo-1 fluorescence ratio. Data is representative of 2 independent experiments (n=4). **I** Normalized baseline Indo-1 ratios in IgE^+^ and IgG1^+^ iGCs and PCs at day 8 of culture. Each dot represents a technical replicate, lines indicate mean. Data were analyzed using a One-way RM ANOVA with Sidak’s multiple comparisons test. **J** Normalized surface IgE-BCR expression in IgE^+^ iGCs compared to IgE^+^ PCs. **K** Normalized surface IgG1 BCR expression in IgG1^+^ iGC compared to IgG1^+^ PC. Cells were cultured in the presence of 500ng/ml anti-Igκ F(ab’)_2_ for the final 2 days of culture. In **J** and **K** cells within the same culture well are connected by a line, data are pooled from 2 separate experiments. Statistical significance (paired t-test) is indicated. **L** Proportions of IgE^+^ PCs (as a % of live lymphocytes) treated with 1μM of the Syk inhibitor BAY 61-3606 (white bars) or vehicle control (black bars) added for the final 2 days of B cell culture (n=10-11, data pooled from 4 separate experiments) bars show mean ± SEM. Statistical significance (unpaired t-test) is indicated. **M** Proportions of active caspase3^+^ sorted IgE^+^ PC treated with 1μM of the Syk inhibitor BAY 61-3606 (open circles) or vehicle control (closed circles) added for 20 hours. Each point represents a biological replicate. **N** Normalized Indo-1 fluorescence ratio in WT (black dots) and *Atp2b1* KO cells at day 8 of culture. **O** Screen validation data showing CRISPR KO of *Atp2b1* using 2 different sgRNAs. Data are pooled from 4 independent experiments (Plots as described for **A**). **P** Proportions of active caspase3^+^ WT (black dots) and *Atp2b1* KO cells at day 8 of culture. In **N** and **P** each dot represents a biological replicate, line indicates mean. Data were analyzed using a Two-way RM ANOVA with Sidak’s multiple comparisons test. **Q** Proportions of IgE^+^ and IgG1^+^ B cells and PCs shown for 10nM thapsigargin treated (white bars) compared to vehicle treated (black bars) cells, where thapsigargin was added for the final 2 days of *in vitro* B cell cultures. Data are pooled from 2 individual experiments (n=3 biological replicates), bars show mean ± SEM. Statistical significance (unpaired t-test) indicated. **R** Proportions of Annexin V^+^ cells in 10nM thapsigargin treated (white bars) compared to vehicle treated (black bars) (as described for **M**). Data is pooled from 3 separate experiments (n=4 biological replicates), bars show mean ± SEM. Statistical significance (unpaired t-test) indicated.

### Mechanisms of calcium-induced IgE^+^ cell death

To understand why IgE^+^ B cells and IgE^+^ PCs appear to be more sensitive to increases in cytosolic calcium, we performed calcium flux analysis on IgE^+^ and IgG1^+^ iGCs and IgE^+^ PCs. We observed that IgE^+^ and IgG1^+^ B cells fluxed calcium to the same extent in response to thapsigargin, ionomycin (Figure 6H) or anti-Igκ F(ab’)_2_ (data not shown), but baseline calcium levels were greater in IgE^+^ cells than IgG1^+^ cells and increased further in IgE^+^ PCs (Figure 6H, colored arrows and 6I), which may be attributable to the continuous low level signaling of the IgE-BCR (Haniuda et al., 2016). We hypothesized that this chronic IgE-BCR signaling may persist after differentiation into PCs, due to elevated surface expression of the IgE-BCR on IgE^+^ PCs compared to IgE^+^ B cells (He et al., 2013; Ramadani et al., 2017), which contrasts with the decreased surface BCR expression on IgG1^+^ PCs (Figure 6J-K). Whilst treatment with the Syk inhibitor (BAY 61-3606) blocked IgE^+^ PC differentiation from culture day 6 as expected (Haniuda et al., 2016) (Figure 6L), culturing sorted IgE^+^ PC in the presence of BAY 61-3606 for 20 hours resulted in decreased proportions of active caspase 3^+^ cells compared to vehicle controls (Figure 6M), suggesting that while Syk is required for PC differentiation, the IgE-BCR continues to signal in IgE^+^ PC and that this is detrimental for their survival.

To discover whether altered cytosolic calcium alone could impact survival of IgE^+^ cells, we tested the roles of the calcium transporters *Atp2b1* and *Atp2a2* identified by the screen. Deletion of *Atp2b1* led to elevated intracellular calcium concentrations in all cell types (Figure 6N), and caused loss of IgE^+^ iGCs, IgE^+^ PCs, and IgG1^+^ PCs, however, increased apoptosis was only observed in IgE^+^ PCs (Figure 6O, P), indicating IgE^+^ PCs are uniquely sensitized to calcium-induced apoptosis. To test the role of *Atp2a2* we cultured cells in the presence of the ATP2A2 inhibitor thapsigargin. This resulted in a substantial loss of IgE^+^ iGCs and IgE^+^ PCs (Figure 6Q), as well as a decrease in the proportions of IgE^+^ and IgG1^+^ PCs when cells were cultured in the presence of thapsigargin for longer timepoints (Supplementary Figure 9D). This was accompanied by increased cell death assessed using Annexin V staining in IgE^+^ iGCs and PCs (Figure 6R, Supplementary Figure 9E). Thus, IgE^+^ PCs and to some extent also IgE^+^ B cells have enhanced basal levels of intracellular calcium and are more sensitive to calcium-induced apoptosis than IgG1^+^ B cells.

### Calcium import into mitochondria causes apoptosis in IgE^+^ PCs

It has been observed previously (Akkaya et al., 2018; Roca et al., 2019) that chronic calcium signaling can result in the increased transfer of calcium from the ER to the mitochondria, causing calcium overload in the mitochondria and triggering of cell death. To understand whether the enhanced sensitivity of IgE^+^ PC to calcium-induced cell death could be explained by this mechanism, we quantified mitochondrial mass and mitochondrial calcium levels. MitoTracker Green staining for mitochondrial mass was slightly lower in IgE^+^ iGCs than in IgG1^+^ iGCs, but markedly reduced in both types of PCs (Supplementary Figure 9F, G). To confirm this finding, we used MitoTracker Orange to quantify mitochondrial mass in *ex vivo* IgE^+^ and IgG1^+^ GC B cells and PCs. We found that IgE^+^ PCs have again decreased mitochondrial mass compared to IgE^+^ GC B cells (Supplementary Figure 9H), however mitochondrial mass was only mildly reduced in IgG1^+^ PCs compared to IgG1^+^ GC B cells (Supplementary Figure 9I). Thus, differentiation of IgE^+^ GC B cells to PCs is associated with markedly reduced mitochondrial mass, consistent with previous findings in un-switched cells (Jang et al., 2015; Lin et al., 2015).

To quantify mitochondrial calcium, we loaded cells with Rhod-2. The intensity of Rhod-2 was lower in IgE^+^ than IgG1^+^ cells, and lower still in PCs (Supplementary Figure 9J), in conjunction with the lower MitoTracker Green staining (Supplementary Figure 9K). To account for the decreased mitochondrial mass in PCs, we normalized the Rhod-2 intensity to MitoTracker Green intensity and found that IgE^+^ PCs had the highest levels of Rhod-2 relative to MitoTracker Green (Supplementary Figure 9L). Blocking mitochondrial calcium uptake by Ru360 had little effect on iGC cells (data not shown), but enhanced proportions of IgE^+^ PCs and to a lesser extent IgG1^+^ PCs differentiated without anti-Igκ F(ab’)_2_ (Supplementary Figure 9M, N). Correspondingly, we observed a decreased percentage of active caspase 3^+^ IgE^+^ PCs that had been cultured in the presence of Ru360 (Supplementary Figure 9O) indicating that the increased proportion of IgE^+^ PCs was due to decreased cell death. The proportion of active caspase 3^+^ IgG1^+^ PCs was not different when comparing cells treated with and without Ru360 (Supplementary Figure 9P). We conclude that the IgE^+^ PCs, which have increased basal levels of cytosolic calcium, undergo calcium-induced apoptosis initiated by calcium entry into mitochondria. The decrease in mitochondrial mass in IgE^+^ PCs both *in vitro* and *ex vivo* may limit buffering of the cytosolic calcium in the cytosol and contribute to the mitochondrial calcium overload.

### Elevated cytosolic calcium drives IgE^+^ PC death through BCL2L11

To further link the sensitivity to calcium-induced cell death and the mitochondrial apoptotic pathway, we tested whether calcium-induced apoptosis depended on BCL2L11. To this end we performed double KO experiments in B cells *in vitro* in which sgRNAs targeting *Bcl2l11* were delivered in a vector co-expressing BFP, and sgRNAs targeting *Atp2b1* were delivered in the mCherry plasmid. In this way, we compared both single KO and double KO populations from within the same culture well (Figure 7A). As seen previously (Figure 5B), *Bcl2l11* single KO resulted in increased proportions of IgE^+^ PCs, which had decreased proportions of active caspase 3^+^ cells (Figure 7B, C). In contrast, the KO of *Atp2b1* resulted in decreased proportions of IgE^+^ PCs and increased proportions of active caspase 3^+^ cells (Figure 7B, C). Within the double KO population, the IgE^+^ PC proportions were increased when compared to WT cells, almost to the level of *Bcl2l11* KO cells, and active caspase 3^+^ proportions were comparable to WT and *Bcl2l11* KO cells (Figure 7B, C), suggesting that elevated cytosolic calcium levels drive apoptosis of IgE^+^ PCs mainly through BCL2L11.

**Figure 7:**
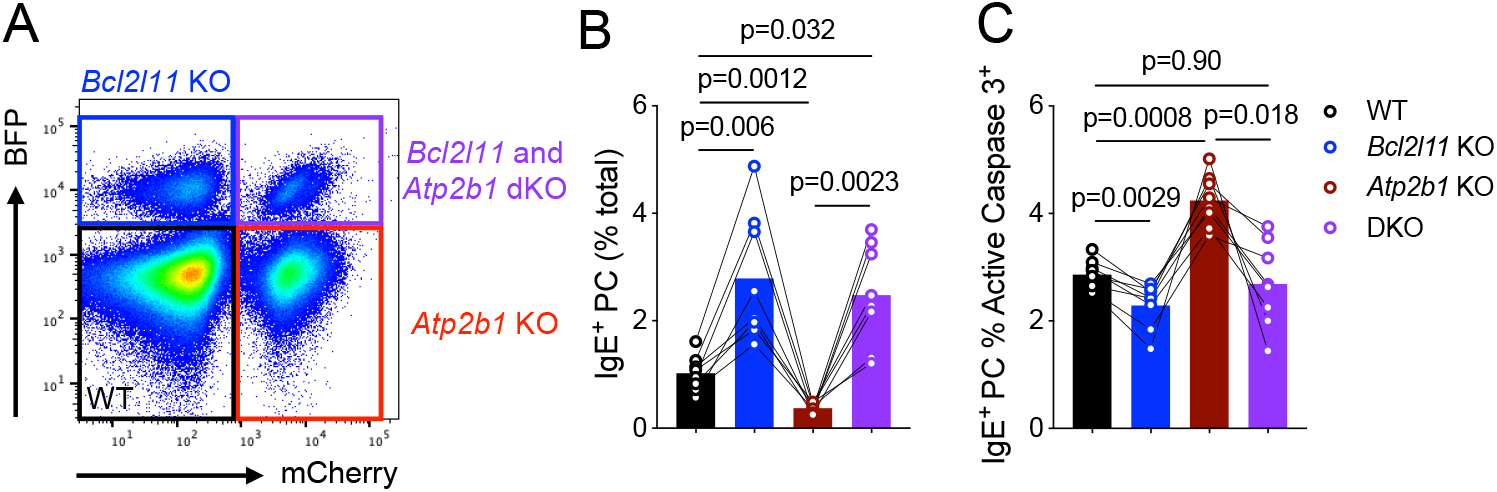
Calcium signaling drives cell death through BCL2L11. **A** Representative FACS plot demonstrating identification of *Bcl2l11* KO (BFP^+^ mCherry^−^), *Atp2b1* KO (BFP^−^ mCherry^+^), WT (BFP^−^ mCherry^−^) and DKO (BFP^+^ mCherry^+^) populations from the same culture well. **B** Proportion of IgE^+^ PC in populations described in **A**. **C** Proportion of active caspase 3^+^ IgE^+^ PC in populations described in **A**. Each bar in **B** and **C** shows mean for 8 replicates, cells within the same culture well (connected by line), data are pooled from 2 separate experiments. Statistical significance (One-way RM ANOVA with Tukey’s multiple comparisons test) shown.

## Discussion

Our study represents a genome-wide identification of genes involved in class-switching and the growth, survival and differentiation of IgG1- and IgE-switched B cells. We confirmed known pathways required for mature B cell survival, such as tonic BCR and BAFFR signaling, as well as key regulators of CD40L and IL-4 dependent proliferation, survival, class-switching and PC differentiation, validating the screening method. These screen data thus present a useful resource to investigate the mechanisms underlying class-switched B cells responses. We demonstrate the utility of this approach by investigating the molecular pathways that control IgE^+^ B cell differentiation and lifespan. The results confirm the importance of antigen-independent IgE-BCR signaling and reveal the downstream mechanisms including identification of previously unreported genes and pathways.

Firstly, we were able to demonstrate a crucial role for the PI3K-mTOR signaling pathway in driving PC differentiation of IgE^+^ B cells. PI3K signaling is a key pathway in immunity and the IgE-BCR has been shown to autonomously induce greater phosphorylation of the co-receptor CD19 and the PI3K p85 subunit than the IgG1-BCR (Haniuda et al., 2016). In B cells, PI3K signaling has been proposed to act upstream of IRF4, a key transcription factor required to initiate PC differentiation (Haniuda et al., 2016; Sciammas et al., 2006), however the molecular mechanisms remain unclear. Our data reveal that in IgE^+^ cells this molecular connection depends on mTOR, which promotes accumulation of IRF4 protein through a post-transcriptional mechanism. Specifically, we found increased IRF4 protein but not mRNA in IgE^+^ B cells and showed that in IgE^+^ PC IRF4 protein levels but not mRNA was diminished by treatment with Rapamycin. As the *Irf4* gene carries a potential 5’ terminal oligopyrimidine (TOP) motif, we hypothesize that PI3K stimulates Irf4 translation via mTOR-mediated phosphorylation of EIF4EBP1 (Thoreen et al., 2012). Although mTOR activity is ultimately required for all PCs, the post-transcriptional regulation of IRF4 levels in IgE^+^ cells illustrates the unique mechanisms that control their differentiation into PCs and suggests that it depends on synergy with factors that drive *Irf4* transcription, such as CD40 and IL-4R signaling.

The second key finding of our screening approach was the identification of the calcium signaling pathway as a negative regulator of PC differentiation and survival. This was unanticipated as calcium signaling has long been thought to promote PC differentiation, particularly in IgG and IgE-switched cells (Engels and Wienands, 2018). The calcium pathway was identified through numerous screen hits that point to a negative role for calcineurin-NFATC3 pathway in PC differentiation, which is particularly strong in IgE^+^ cells. This suggests that B cell intrinsic roles contribute to the previously identified role for NFAT proteins in inhibiting IgE and IgG1 production (Peng et al., 2001), their association with autoimmunity and allergy in GWAS studies(Ferreira et al., 2017), and enhanced allergic responses associated with elevated serum IgE levels in transplant patients treated with cyclosporin A or tacrolimus, both of which inhibit calcineurin (Asante-Korang et al., 1996; Kawamura et al., 1997). Furthermore a recent forward genetic analysis in mice identified mutations in *Plcg2* and *Syk*, which resulted in augmentation of the IgE response to papain immunization (SoRelle et al., 2020). Human mutations in *PLCG2*, which attenuate calcium signaling in B cells (Wang et al., 2014) also lead to enhanced allergen-specific IgE levels in the context of phospholipase-associated antibody deficiency and immune dysregulation (Milner, 2020). Calcineurin may inhibit PC differentiation transcriptionally through the activity of NFATC3 and its target genes. However, since deletion of *Ppp3r1* had a stronger phenotype in IgE^+^ cells than in IgG1^+^ cells, a phenotype that differed from that of *Nfatc3* deletion, it is possible that other PPP3R1 targets are important in regulation of IgE^+^ responses, such as *Crtc2*. Despite this, we are intrigued by the non-redundant function of NFATC3 in this process, and postulate that this may in part be accounted for by the differential regulation of NFAT family members by discrete calcium signals (Kar et al., 2016). Indeed, NFATC3 is readily exported from the nucleus, and therefore requires sustained calcium signals to maintain nuclear localization. This aligns with the inhibitory role for the IP_3_R-2 in PCs, which has been shown to be the most sensitive IP_3_R to IP_3_ levels and to mediate long-term calcium oscillations downstream of the BCR (Miyakawa et al., 1999), such as those required for NFATC3 nuclear localization.

Apoptosis has been previously suggested to limit the expansion of IgE^+^ B cells (Haniuda et al., 2016; Oberndorfer et al., 2006), however the importance of this pathway remains controversial (Yang et al., 2016), in part because a molecular mechanism remains to be elucidated. Our data suggest that IgE^+^ B cell and PC death is regulated by BCR-induced ER calcium release and a balance of proapoptotic (BCL2L11 and BAX) and anti-apoptotic (BCL2A1D/B) proteins. This finding is mirrored by the altered expression levels of apoptotic proteins in mouse and human IgE^+^ B cells (Ramadani et al., 2019). As has been observed previously (Haniuda et al., 2016), IgE^+^ B cells exhibit increased basal calcium levels, which we show is further increased in IgE^+^ PCs. Notably, the intracellular calcium levels in IgE^+^ PCs depend on SYK, indicating that they are driven by ongoing IgE-BCR signaling. Consistently, inhibition of SYK prevents the apoptosis of isolated IgE^+^ PCs, indicating that IgE-BCR signaling regulates their survival after differentiation. It is interesting to speculate that this mechanisms arises from elevated surface expression of the IgE-BCR on these cells (He et al., 2013; Ramadani et al., 2017).

Mechanistically, previous observations indicate that chronic elevation of intracellular calcium downstream of the BCR leads to mitochondrial disfunction and cell death (Akkaya et al., 2018). Mitochondrial dysfunction caused by excessive entry of calcium into mitochondria is itself regulated by pro- and anti-apoptotic proteins. For example, it has been shown in the context of macrophages that BAX can promote calcium flow from the ER into mitochondria resulting in cell death (Roca et al., 2019). Conversely, BCL2 has been shown to reduce mitochondrial Ca^2+^ uptake by blocking IP_3_-mediated Ca^2+^ release from the ER through inhibition of IP_3_Rs in a T cell line (Chen et al., 2004). We observed decreased mitochondrial mass upon PC differentiation, which was augmented in IgE^+^ PCs. This is in line with the proposed transcriptional regulation of mitochondrial metabolism and mass by IRF4 in early plasma cells (Low et al., 2019). The decreased mitochondrial mass in IgE^+^ PCs also fits with the notion that reductions in mitochondrial mass correlate with a predisposition of B cells to differentiate into PC (Jang et al., 2015). Overall, this suggests that impairment of mitochondrial calcium buffering alongside with increased cytosolic release from ER is a key mechanism by which the lifespan of IgE^+^ cells is limited. Indeed, although the role of apoptosis in IgE^+^ GC B cells has been controversial, IgE^+^ PCs have a high level of apoptosis *in vivo* and their numbers are specifically increased by overexpression of the anti-apoptotic protein BCL2 (Yang et al., 2012, 2020). Thus, BCR apoptotic signaling in IgE^+^ PCs could be responsible for their short lifespan. As further remodeling of mitochondria has been suggested in long-lived plasma cells (Low et al., 2019), it will be interesting to understand the importance of calcium signaling in regulating survival of the recently identified long-lived IgE^+^ PCs generated after chronic allergen exposure (Asrat et al., 2020).

Overall, our genome-wide CRISPR screening indicate that the chronic dynamics of IgE-BCR signaling translates into a distinct quality of calcium signaling in IgE^+^ B cells and is responsible for curtailing IgE^+^ PC differentiation and survival. The stringent control of the IgE^+^ cell fates help to explain the self-limiting nature of IgE responses and suggests that dysregulation of this pathway may contribute to allergy.

## Acknowledgements

This work was supported by the Francis Crick Institute which receives its core funding from Cancer Research UK (FC001185), the UK Medical Research Council (FC001185), and the Wellcome Trust (FC001185).

This research was funded in whole, or in part, by the Wellcome Trust (Grant number FC001185). For the purpose of Open Access, the author has applied a CC BY public copyright license to any Author Accepted Manuscript version arising from this submission.

We thank the Francis Crick Institute Science Technology Platforms for their help with cell-sorting and next generation sequencing, and Harshil Patel from the Bioinformatics and Biostatistics Science Technology Platform at The Francis Crick Institute for performing the bioinformatics analysis of the RNAseq data.

We also thank Daisuke Kitamura for the 40LB cells, and Lars Nitschke and Richard Cornall for sharing the *Grb2*^fl/fl^*Cd79a*^cre/+^ mice. We also thank Edina Schweighoffer and Victor Tybulewicz for providing the Brie CRISPR library and the Syk inhibitor BAY 61-3606.

## Author contributions

R.N. designed and conducted the experiments and analyzed the data, P.T. designed the experiments analyzed the data and supervised the research. R.N. and P.T. wrote the paper.

## Declaration of Interests

The authors declare no competing interests.

---

**Supplementary Figure 1: Production and characterization of Cherry Brie library**

**A** Schematic depicting Cherry Brie library plasmid map. **B**, **C** Histograms depicting the distribution of normalized read counts per sgRNA within the original Brie library (**B**) and in the new Cherry Brie library (**C**). **D**-**F** Histograms depicting the distribution of normalized read count per sgRNA from pooled CRISPR screens in IgG1^+^ iGC cells (**D**), IgE^+^ iGC cells (**E**), IgE^+^ PCs (**F**). **G**-**I** Histograms depicting CRISPR scores for whole library (All, black line), common essential genes (Wang et al., 2015) (Essential, blue bars) and non-targeting control guides (Controls, Pink bars) for IgG1^+^ iGC CRISPR screen (**G**), IgE^+^ iGC CRISPR screen (**H**), IgE^+^ PC CRISPR screen (**I**). **J**-**L** Box and whisker plots showing distribution of CRISPR scores for whole library (All, black), common essential genes (Wang et al., 2015) (Essential, blue) and non-targeting control guides (Controls, pink) for IgG1^+^ iGC CRISPR screen (**J**), IgE^+^ iGC CRISPR screen (**K**), IgE^+^ PC CRISPR screen (**L**). For IgG1^+^ iGC CRISPR screen, data is pooled from three technical replicates, for IgE^+^ iGC CRISPR screen, data is pooled from two technical replicates.

**Supplementary Figure 2: RNAseq of *in vitro* B cell culture**

**A** Schematic depicting RNAseq workflow. **B** Principal component analysis (PCA) plot indicating distinct gene expression profiles of sorted and bulk-day 6 populations. Each point represents a single biological replicate. **C**-**J** Transcript abundance in sorted populations (as indicated in **A)** of Bcl6 (**C**), Pax5 (**D**), Prdm1 (**E**), Irf4 (**F**), Xbp1 (**G**), Ighe (**H**), Ighg1 (**I**), Ighm (**J**). Each bar represents 3 biological replicates, bars show mean ± SEM.

**Supplementary Figure 3: Impact of adding soluble anti-Igκ as a surrogate antigen into B cell cultures**

PC differentiation in B cell cultures following 6 days of culture on 40LB feeder cells in the presence of IL-4, followed by 2 days in culture with and without soluble anti-Igκ F(ab’)_2_. PCs were gated as CD138^+^ B220^lo^ cells. **A** Effect of soluble anti-Igκ F(ab’)_2_ concentration on proportions of IgE^+^ PCs shown as percentage of total IgE^+^ cells. **B** Representative FACS plots indicating percentage of PCs in cultures with and without soluble anti-Igκ F(ab’)_2_ within IgE^+^ gate. Numbers on plots indicate the percentage of cells within the CD138^+^ B220^lo^ gate. **C** Effect of soluble anti-Igκ F(ab’)_2_ concentration on proportions of IgG1^+^ PCs shown as percentage of total IgG1^+^ cells. **D** Representative FACS plots indicating percentage of PC in cultures with and without soluble anti-Igκ F(ab’)_2_ within IgG1^+^ gate. Numbers on plots indicate the percentage of cells within the CD138^+^ B220^lo^ gate. For **A** and **C,** bars are indicative of a single replicate (n=1). For **C** and **D** FACS plots are representative of 13 replicates.

**Supplementary Figure 4: Identification of common expected essential genes**

**A** Comparative dot plot indicating gene CRISPR scores in the IgE^+^ iGC screen and IgG1^+^ iGC screens. Expected common essential genes (blue points) as indicated in volcano plots in **Figure 1B, C**. Common negative regulators shown as red points. **B** Comparative dot plot indicating genes essential in the IgE^+^ iGC screen and IgE^+^ PC screens. Expected common essential genes (blue points) as indicated in volcano plots in **Figure 1C, D**. Common negative regulators shown as red points.

**Supplementary Figure 5: CRISPR screens identify Morc3 as a novel regulator of B cell development**

**A** Comparative dot plot indicating gene CRISPR scores of Morc3 (shown as a red dot) in the IgE^+^ iGC screen and IgG1^+^ iGC screens. **B** Validation of *Morc3* essentiality by CRISPR KO effect on IgE^+^ and IgG1^+^ cell numbers. Bars show mean ± SEM of proportions of mCherry^+^ IgE^+^ and IgG1^+^ cells at day 8 of B cell culture relative to proportions of mCherry^+^ B cells at day 3 of B cell culture. Data are pooled from KO effects of 2 sgRNAs per gene (n=2). **C** Schematic of CRISPR-Cas9 chimera generation. **D**-**I** Targeting of *Morc3* and *Cd4* as a control. Data show proportions of Transitional 1 B cells (**D**), Transitional 2 B cells (**E**), Transitional 3 B cells (**F**), Follicular B cells (**G**) Marginal zone B cells (**H**) and CD4 T cells (**I**) in the spleen from CRISPR Cas9 chimeras comparing WT (GFP^+^ mCherry^−^) black bars and KO (GFP^+^ mCherry^+^) red bars. For **D**-**I** data represent mean ± SEM of 5 mice from 1 experiment. Statistical significance (Two-way RM ANOVA with Sidak’s multiple comparisons test) is indicated.

**Supplementary Figure 6: Validation of *Irf4* and *Prdm1* as central regulators of IgE^+^ PC differentiation**

**A** Screen validation data showing KO of *Irf4* using 2 different sgRNAs (shown as open and closed circles). Proportions of IgE^+^ cells (left graph) and IgE^+^ PC (right graph) shown for WT (GFP^+^ mCherry^−^) and KO (GFP^+^ mCherry^+^) cells within the same culture well (connected by line). Data are pooled from 5 separate experiments. **B** Screen validation data showing KO of *Prdm1* using 2 different sgRNAs. (Plots as described for **A**). Data pooled from 2 separate experiments. Statistical significance for **A** and **B** in paired t-tests is indicated.

**Supplementary figure 7: mTOR regulation of PC differentiation**

**A** Proportions of iGCs and PCs (as a % of live lymphocytes) with 100nM Rapamycin (white bars) or vehicle control (black bars) added for the final 2 days of B cell culture. Data are pooled from 2 separate experiments showing mean ± SEM (n=3 biological replicates). Statistical significance (unpaired t-test) is indicated. **B**, **C** Histograms showing expression of phospho-mTOR (Ser2448) in IgE^+^ or IgG1^+^ iGCs (grey filled histogram) compared to IgE^+^ PCs or IgG1^+^ PCs respectively (black line) and a secondary antibody only staining control (black dotted line) (**B**), phospho-S6 (Ser235/236) in IgE^+^ or IgG1^+^ iGCs (grey filled histogram) compared to IgE^+^ PCs or IgG1^+^ PCs respectively (black line) and a secondary antibody only staining control (black dotted line) (**C**) at day 8 of *in vitro* cultures without surrogate antigen. Data in **B** and **C** are representative of 3 biological replicates from 2 separate experiments. **D**, **E** FPKM values from RNAseq data indicating transcript abundance in IgE^+^ vs IgG1^+^ iGC and PC of *Irf4* (**D**), *Prdm1* (**E**). Each dot represents a biological replicate, line indicates the mean. Statistical significance (unpaired t-test) indicated. **F**, **G** Median fluorescence intensity (MFI) of IRF4 (**F**), PRDM1 (**G**) in flow cytometry on IgE^+^ vs IgG1^+^ iGCs and PCs. Each dot represents a technical replicate, line indicates mean. Data are representative of 3 independent experiments, statistical significance (unpaired t-test) indicated.

**Supplementary Figure 8: Regulation of IgE^+^ B cell responses by genes involved in calcium signaling**

**A** Screen validation data showing CRISPR KO of *Grb2* using 2 different sgRNAs (shown as open and closed circles). Conditions where 500ng/ml anti-Igκ F(ab’)_2_ was added for the final 2 days of culture are shown in blue. Proportions of IgE^+^ cells, IgE^+^ PCs, IgG1^+^ cells, and IgG1^+^ PCs (graphs from left to right) shown for WT (GFP^+^ mCherry^−^) and KO (GFP^+^ mCherry^+^) cells within the same culture well (connected by line), data are pooled from 4 separate experiments. Data in **A** were analyzed using paired t-tests. Significance values are indicated for cultures with (blue) and without (black) addition of anti-Igκ F(ab’)_2_. **B**, **C** Proportion of IgE^+^ GC B cells (**B**) and IgG1^+^ GC B cells (**C**) at 7 days after immunization in draining LN of *Grb2*^fl/fl^*Cd79a*^Cre/+^ and *Grb2*^fl/fl^ littermate controls immunized subcutaneously with NP-CGG in Alum. In **B**-**C** each dot represents a biological replicate, line represents mean. Data were pooled from 2 independent experiments and analyzed using an unpaired t-test. **D** Screen validation data showing KO of *Itpr2* using 2 different sgRNAs, data are pooled from 2 separate experiments (plots as described for **A**). **E** Screen validation data showing CRISPR KO of *Itpr3* using 2 different sgRNAs, data are pooled from 2 separate experiments (plots as described for **A**). **F** Screen validation data showing KO of *Crtc2* using 2 different sgRNAs. Data are pooled from 2 separate experiments. (Plots as described for **A**). **G** Screen validation data showing KO of *Plcg2* using 2 different sgRNAs. Data are pooled from 6 separate experiments. (Plots as described for **A**). **H-K** Proportions of GC B cells (**H**), IgE^+^ GC B cells (**I**), IgG1^+^ GC B cells (**J**), and PCs (**K**) in draining LN from CRISPR Cas9 chimeras comparing WT (GFP^+^ mCherry^−^) black bars and KO (GFP^+^ mCherry^+^) red bars. For **H**-**K** data are pooled from 2 separate experiments, each bar is representative of 8 mice, bars represent mean ± SEM. Statistical significance (Two-way RM ANOVA with Sidak’s multiple comparisons test) is indicated.

**Supplementary Figure 9: Mitochondrial dysregulation in PC**

**A** Proportion of active caspase-3^+^ IgE^+^ and IgG1^+^ cells and PCs from day 8 of *in vitro* culture, comparing WT (black bars, open circles) to *Plcg2* KO (white bars, closed circles). Cells within the same culture well are connected by a line, data are pooled from 2 separate experiments. Statistical significance (Two-way RM ANOVA with multiple comparisons test) indicated. **B** Screen validation data showing KO of *Itpkb* using 2 different sgRNAs (shown as open and closed circles). Conditions where 500 ng/ml anti-Igκ F(ab’)_2_ was added for the final 2 days of culture are shown in blue. Proportions of IgE^+^ cells, IgE^+^ PCs, IgG1^+^ cells and IgG1^+^ PCs (graphs from left to right) shown for WT (GFP^+^ mCherry^−^) and KO (GFP^+^ mCherry^+^) cells within the same culture well (connected by line). Data are pooled from 6 separate experiments. **C** Proportions of Annexin V^+^ cells in WT (black bars) and *Itpkb* KO (white bars) data is pooled from two experiments (n=6). Bars show mean ± SEM. Statistical significance (Two-way ANOVA with Sidak’s multiple comparisons test) indicated. **D** Bar graph showing the proportions of total IgE^+^ and IgG1^+^ cells and PC (as a % of live lymphocytes) with 10nM thapsigargin (white bars) or vehicle control (black bars) added for the final 5 days of B cell culture. Data are pooled from 2 separate experiments, (n=3) bars show mean ± SEM. **E** Bar graph showing the proportions of Annexin V^+^ cells in total IgE^+^ and IgG1^+^ cells and PC with 10nM thapsigargin (white bars) or vehicle control (black bars) added for the final 5 days of B cell culture. Data is pooled from 3 separate experiments (n=4), bars show mean ± SEM. Data in **D** and **E** were analyzed using unpaired t-tests, statistical significance is indicated. **F** MitoTracker Green normalized MFI in IgE^+^ and IgG1^+^ iGC and PC populations. Each dot represents a technical replicate, line indicates mean, data are pooled from 2 independent experiments. **G** Fluorescence intensity of MitoTracker Green staining per mitochondria using fluorescence microscopy. Each dot represents a single mitochondrion, bars indicate mean. Data in **F and G** were analyzed using a One-way ANOVA with Tukey’s multiple comparisons test, statistical significance is indicated. **H, I** MitoTracker Orange normalized MFI in *ex vivo* IgE^+^ (**H**) and IgG1^+^ (**I**) GC B cells and PCs from mice 7 days after immunization with NP-CGG subcutaneously. Each dot represents a mouse, line indicates mean, data are pooled from 2 independent experiments. Data were analyzed using an unpaired t-test. **J** Rhod-2 normalized median fluorescence intensity (MFI) in IgE^+^ and IgG1^+^ iGC and PC populations. Each dot represents a technical replicate, line indicates mean, data pooled from 3 independent experiments, data were analyzed using a One-way ANOVA with Tukey’s multiple comparisons test. **K** Representative flow cytometry contour plots indicating Rhod-2 vs MitoTracker Green staining in total IgE^+^ shown in red, overlaid with total IgG1^+^ blue (left plot), IgE^+^ (grey), overlaid with IgE^+^ PCs shown in red (middle plot), and IgG1^+^ (grey) overlaid with IgG1^+^ PCs shown in blue (left plot), with adjunct histograms. Data are representative of 2 experiments (n=3). **L** Rhod-2 MFI normalized to MitoTracker Green MFI in each cell type. Data is representative of 3 biological replicates from 2 independent experiments. **M**, **N** Proportions of IgE^+^ PCs (**M**) IgG1^+^ PCs (**N**) with and without 10μM Ru360 for the final 24 hours of culture. Conditions where 500 ng/ml anti-Igκ F(ab’)_2_ was added for the final 2 days of culture are shown in blue. Data are pooled from 3 separate experiments, (n=8). Points indicate mean proportions ± SEM. **O**, **P** Proportions of Active Caspase 3^+^ cells in vehicle and 10μM Ru360 treated IgE^+^ PC (**O**) IgG1^+^ PC (**P**). Conditions where 500 ng/ml anti-Igκ F(ab’)_2_ was added for the final 2 days of culture are indicated. Data is from one experiment (n=4). Data in **M**-**P** were analyzed using paired t-tests. Significance values are indicated for cultures with (blue) and without (black) addition of anti-Igκ F(ab’)_2_.

## Materials and Methods

### Reagents, antibodies and oligonucleotides

This information is provided in Resources Table.

### Mouse strains and animal procedures

Mice used in this study were rederived onto the C57BL/6 background at the Francis Crick Institute. Cas9-GFP mice used were Gt(ROSA)26Sor^tm1.1(CAG-cas9*,-EGFP)Fezh^ (MGI allele ID 5583838). *Grb2*^fl/fl^ mice used were B6.C(Cg)-*Grb2*^tm1.1Lnit^/J (MGI allele ID 95805). *Cd79a*^cre/+^ mice used were C(Cg)-Cd79a^tm1(cre)Reth^/EhobJ (MGI allele ID 101774). Tissues were taken from mice aged between 8-15 weeks. All experiments were approved by the Francis Crick Institute Ethical Review Panel and the UK Home Office.

### Flow cytometry

Single cell suspensions were blocked with anti-CD16/32 for 20 minutes and incubated with Fixable Viability Dye-ef780 (ThermoFisher) for 20 minutes. Cells were subsequently stained with appropriate antibodies for 20 minutes on ice. The following stains and antibodies (for further details see Resources Table) were used for immunophenotyping: B220 (RA3-6B2), CD138 (281-2), CD21/35 (eBio4E3), CD23 (B3B4), CD38 (90), CD4 (RM4-5), CD93 (AA4.1), CD95 (Jo2), IgD (11-26c), IgG1 (RMG1-1 or X56), IgM (R6-60.2), IgE (RME-1), TCRβ (H57-597). Cells were analyzed using an LSR-Fortessa flow cytometer. To measure apoptosis, cells were stained using the Annexin V apoptosis detection kit (ThermoFisher) following manufacturer’s instructions. Intracellular staining using antibodies detecting Active Caspase-3 (C92-605) and BCL2L11 (Y36) was performed using BD Cytofix/Cytoperm (BD Biosciences) or eBioscience FoxP3/Transcription Factor Staining Buffer Set (ThermoFisher). Intracellular staining using antibodies to detect IRF4 (3E4) and PRDM1 (5E7) was performed using True-Nuclear Transcription Factor Buffer Set (Biolegend). For mitochondrial stains, cells were loaded for 30 minutes at 37°C in the dark in HBSS with 500nM Mitotracker^TM^ Green FM (ThermoFisher), 500nM MitoTracker^TM^ Orange CMTM Ros (ThermoFisher), 100nM TMRM (AAT Bioquest) or 5μM Rhod-2 AM (ThermoFisher). Rhod-2 AM was loaded in the presence of 0.02% Pluronic F127 (ThermoFisher). Intracellular staining for phospho-S6 and phospho-mTOR (both from Cell Signaling Technology) using eBioscience FoxP3/Transcription Factor Staining Buffer Set (ThermoFisher).

### Calcium flux analysis

IgG1^+^ and IgE^+^ B cells from the *in vitro* culture (8x×10^6^ cells) were loaded for 30 minutes at 37°C in the dark with 4.5μM Indo-1 AM (ThermoFisher) in the presence of 0.02% Pluronic F127 (ThermoFisher). Cells were then stained with surface antibodies or Fab fragments against IgE (made in-house) and IgG (Jackson ImmunoResearch Europe Ltd) at RT and resuspended in HBSS. Cells were pre-warmed to 37°C before analysis. After 60s of acquisition, cells were stimulated with 1μM Thapsigargin (Sigma-Aldrich). Samples were acquired for 8 minutes, before addition of 5μg/ml Ionomycin (Sigma-Aldrich). Indo-1 fluorescence emission was measured using which a 355nm laser and 450/50 (Indo-1 violet) and 530/50 (Indo-1 blue) filter sets on a BD LSR-Fortessa flow cytometer. The ratio was calculated as Indo-1 violet/Indo-1 blue.

### B cell purification and sorting

Naïve primary B cells were isolated from C57BL/6 or Cas9-GFP (described above) mice. B cells from the spleen were isolated by negative selection using ACK lysis (made in house) and CD43 microbeads (Miltenyi Biotech), following manufacturer’s instructions. To purify cultured B cells from 40LB feeders, the Feeder Removal Kit, mouse (Miltenyi Biotech) was used, following manufacturer’s instructions. To purify IgE^+^ and IgG1^+^ cells for RNAseq or for CRISPR-Cas9 screens, cells were sorted according to the staining strategy as described below using a BD FACS Aria III or a BD FACS Aria Fusion. Sorted IgE^+^ PC for culture with BAY 61-3606 were magnetically pre-enriched using negative selection and were stained using a Fab fragment against IgE (made in house) and anti-CD138.

### 40LB cultures

Naïve primary mouse B cells were isolated as described above. 5×10^5^ B cells were grown on 5×10^5^ 40LB feeder cells in each well of a 6 well plate. Cells were cultured in RPMI media (Sigma) supplemented with 10% heat-inactivated fetal bovine serum (FBS) (ThermoFisher), 100μM non-essential amino acids (ThermoFisher), 2mM L-Glutamine (ThermoFisher), 50μM 2-Mercaptoethanol (ThermoFisher) and Penicillin-Streptomycin (GE Healthcare Life Sciences). 100nm IL-4 (Peprotech) was added into the culture media. Cells were cultured on the same feeders for 3 days before being moved onto new feeders and put into fresh media with IL-4 for a further 3 days. After 6 days culture in the presence of 40LB cells, the B cells were isolated from the feeders using the Feeder Removal Kit, mouse (Miltenyi Biotech) following manufacturer’s instructions. B cells were then cultured for a further 2 days in RPMI (as described above) or supplemented with 500ng/ml Igκ (Southern Biotech). Where indicated the following reagents were added into *in vitro* cultures: 100nM Rapamycin, 10nM Thapsigargin, 10μM Ru360 (all Sigma-Aldrich), 1μM BAY 61-3606 (Calbiochem).

### RNAseq

B cells were isolated from the spleens of 3 C57BL/6 female mice (aged 11 weeks) using CD43 Microbeads (Miltenyi Biotech) as described previously. For each biological replicate, B cells were cultured on 2 6-well plates of 40LB feeders as described previously, in the presence of 100ng/ml IL-4 (Peprotech). B cells were removed from feeders as described previously and cultured in RPMI without stimulation for 2 days prior to sorting. Cells were sorted as follows: IgE iGC (IgE^+^, CD138^−^), IgG1 iGC (IgG1^+^, CD138^−^), IgE PC (IgE^+^, B220^lo^, CD138^−^), IgG1 PC (IgG1^+^, B220^lo^, CD138^−^). Sorted cells were immediately lysed in buffer RLT and samples were homogenized by pipetting back and forth through a 25G needle and frozen on dry ice. Samples were stored at −80°C until processed. Bulk d6 samples were taken from a separate B cell culture, as described above, using B cells isolated from the spleens of 3 C57BL/6 female mice (aged 8 weeks). B cells were removed from feeders using Feeder Removal Microbeads, mouse (Miltenyi Biotech) and cells were lysed and frozen as described above.

RNA was isolated from cells as using the RNeasy micro kit (Qiagen) for PC samples or the RNeasy mini kit (Qiagen) for iGC and bulk d6 samples as per manufacturer’s instructions. RNA QC was performed by the Crick HTS (high-throughput sequencing) core facility using a Bioanalyzer (Agilent), all samples had a RIN score of 10. Stranded mRNA libraries were prepared in the Crick HTS facility using a KAPA mRNA HyperPrep Kit (Roche) and sequenced on the Illumina HiSeq 4000 platform (76bp single-end sequencing).

### Bioinformatics for RNAseq

RNA sequencing was carried out on the Illumina HiSeq 4000 platform and typically generated ∼44 million 76bp, strand-specific, single-end reads per sample. Adapter trimming was performed with cutadapt (version 1.9.1) (Martin, 2011) with parameters “--minimum-length=25 --quality-cutoff=20 -a AGATCGGAAGAGC”. The RSEM package (version 1.3.0) (Li and Dewey, 2011) in conjunction with the STAR alignment algorithm (version 2.5.2a) (Dobin et al., 2013) was used for the mapping and subsequent gene-level counting of the sequenced reads with respect to all *M. musculus* genes downloaded from the Ensembl genome browser (assembly GRCm38, release 89) (Kersey et al., 2016). The parameters used were “--star-output-genome-bam --star-gzipped-read-file --forward-prob 0”, and all other parameters were kept as default.

Differential expression analysis was performed with the DESeq2 package (version 1.12.3) (Love et al., 2014) within the R programming environment (version 3.3.1) (R Core team, 2015). An adjusted p-value of ≤0.01 was used as the significance threshold for the identification of differentially expressed genes.

The RNAseq data are available in the Gene Expression Omnibus (GEO) database (http://www.ncbi.nlm.nih.gov/gds) under the accession number GSE166853.

### Production of Mouse sgRNA Cherry Brie Library

LentiGuide-Cherry plasmid was produced by restriction digest cloning, replacing the Puromycin selection cassette in LentiGuide-Puro (Addgene #52963) with the sequence encoding mCherry. The Cherry Brie pooled CRISPR library was obtained by Gibson assembly cloning to place the sgRNA sequences form the Mouse Brie CRISPR knockout pooled library (a gift from David Root and John Doench (Addgene #73633)) into the LentiGuide-Cherry plasmid (Supplementary Figure 1A). The cloned library was expanded in Electrocompetent E.coli (Lucigen) and sequence complexity was analyzed using next generation sequencing using an Illumina MiSeq. Pooled lentiviral libraries were subsequently produced as described below.

### Lentivirus Production

Replication incompetent lentiviruses were produced by co-transfecting 5×10^6^ HEK293T cells with 10μg vector plasmid LentiGuide-Cherry or Cherry Brie, 5μg envelope plasmid pMD2.G (A gift from Didier Trono (Addgene #12259)) or pHIT123 (Soneoka et al., 1995) and 7.5μg packaging plasmid psPAX2 (A gift from Didier Trono (Addgene #12260)) in opti-MEM media (ThermoFisher) using transit-LT1 transfection reagent (Mirus bio LLC). Lentivirus was harvested 48 and 72 hours post-transfection and concentrated by ultracentrifugation. Lentivirus production was scaled up where necessary for production of lentiviral library. Genomic DNA was isolated from HEK293T cells using DNeasy Blood and Tissue Kit (Qiagen) following manufacturer’s instructions.

### CRISPR screens and analysis

Naïve primary mouse B cells were isolated from Cas9-GFP spleens as described above. 50×10^6^ primary B cells were spin-infected for 90mins at 2300rpm in the presence of 16μg/ml polybrene (Millipore) with pooled whole-genome Cherry-Brie lentiviral CRISPR libraries at an MOI of 0.3 to obtain 200-fold coverage of the library. Transduced B cells were cultured on 40LB feeder cells as described previously, B cells were removed from feeders and cultured in RPMI without stimulation for 2 days prior to sorting. To isolate IgE^+^ B cells from CRISPR-Cas9 screen cultures, IgE^+^ cells were first enriched by negative selection, removing IgM^+^ and IgG1^+^ cells using biotinylated antibodies and streptavidin conjugated microbeads (Miltenyi Biotech). Successfully transduced, mCherry^+^ cells were sorted as described for RNAseq experiments. Where necessary cells were pooled from multiple sorts to ensure library coverage, with a minimum of 3×10^6^ sorted cells used per library. Genomic DNA was isolated from sorted cells using the DNeasy Blood and Tissue Kit (Qiagen) following manufacturer’s instructions. The integrated viral genome was amplified using a nested PCR (primers as described in Resources Table). Libraries were sequenced using an Illumina MiSeq. Raw reads in demultiplexed FASTQ files were trimmed to identify sgRNA sequences and aligned to the Brie library sequences using Bowtie. The frequency of aligned read counts for each sgRNA was calculated using MATLAB (Mathworks). CRISPR scores calculation and statistical analysis was performed using MAGeCK (Li et al., 2014).

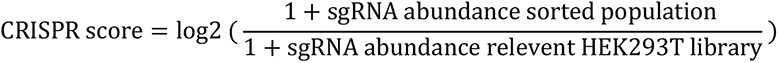

To compare CRISPR scores for the different populations, which were analyzed in separate experiments, CRISPR scores in each population were normalized using:

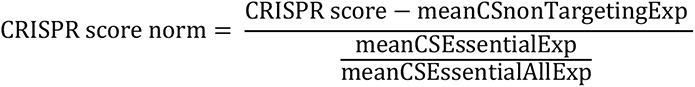

Where meanCSnonTargetingExp is the mean CRISPR score of the non-targeting guides in this experiment, meanCSEssentialExp is the mean CRISPR score of the 187 essential genes (Supplementary Figure 1) in this experiment and meanCSEssentialAllExp is the mean CRISPR score of the 187 essential genes in all experiments.

For IgE^+^ iGC specific gene lists, normalized CRISPR score data was filtered using the following cut-offs: IgE^+^ specific essential genes:

IgE^+^ iGC CRISPR score ≤ −1.8, FDR ≤ 0.25

-0.5 ≤ IgG1^+^ iGC CRISPR score ≤ 0.5

IgE^+^ specific negative regulators:

IgE^+^ iGC CRISPR score ≥ 0.95, FDR ≤ 0.25

-0.5 ≤ IgG1^+^ iGC CRISPR score ≤ 0.5

IgE^+^ PC specific gene lists, CRISPR score data was filtered using the following cut-offs:

IgE^+^ PC specific essential genes:

IgE^+^ PC CRISPR score ≤ −1.5, FDR ≤ 0.25

-1 ≤ IgE^+^ iGC CRISPR score ≤ 0.5

IgE^+^ PC specific negative regulators:

IgE^+^ PC CRISPR score ≥ 0.9, FDR ≤ 0.25 in IgE^+^ PC screen

-0.5 ≤ IgE^+^ iGC CRISPR score ≤ 0.5

To minimize noise in these lists, genes with an average FPKM (in RNAseq analysis) < 10 in all populations were removed.

MLE analysis was performed using MAGeCK by comparing sgRNA counts of the IgE^+^ iGCs and the IgG^+^ iGC or the IgE^+^ PCs and the IgE^+^ iGCs.

### CRISPR-mediated gene disruption

CRISPR sgRNA sequences were designed using the Broad Institute sgRNA Designer. Forward and reverse oligonucleotide sequences including the guide sequence were synthesized, phosphorylated, annealed and individually cloned into lentiGuide-Cherry. Lentivirus was produced and B cells or HSCs were spin-infected as described above. For gene targeting in HSCs, the pMD2.G envelope was used, and for primary B cells, the pHIT123 envelope was used. HSCs were isolated using CD117 beads (Miltenyi Biotech) following manufacturer’s instructions. Spin-infected HSCs were cultured overnight in StemSpan media (StemCell Technologies) in the presence of 100ng/ml mSCF, 6ng/ml IL-3, 10ng/ml hIL-6, 20ng/ml TPO, 60ng/ml Flt3L (all Peprotech).

### Lentiviral chimeras

For bone marrow chimeras RAG-1 KO recipient mice (B6.129S7-*Rag1^tm1Mom^/J*) MGI allele ID 1857241 or Rag1^tm1Bal^ MGI allele ID 2448994 (Spanopoulou et al., 1994)) were sub-lethally irradiated (5Gy) and reconstituted with 1-1.5×10^6^ lentivirally transduced (described above) cKit^+^ donor Cas9-GFP or *Cd79*^Cre/+^ LSL-Cas9-GFP hematopoietic stem cells (HSCs) by intra-venous injection. Reconstituted mice were fed 0.2mg/ml Baytril (Enrofloxacin) in their drinking water for 4 weeks post-reconstitution. Mice were bled to ensure successful reconstitution and were immunized at 8-11 weeks. Mice were immunized sub-cutaneously with 50μl NP-CGG (2B Scientific) (1mg/ml in PBS) mixed 1:1 with Imject Alum Adjuvant (ThermoFisher) in the upper and lower flank to target the brachial, axillary and inguinal draining LNs. Mouse tissues were analyzed by flow cytometry 1 week post-immunization.

### qPCR

RNA was isolated from sorted cells using RNeasy micro kit (Qiagen) or RNeasy mini kit (Qiagen), depending on cell number, following manufacturer’s instructions. cDNA was prepared using the QuantiTect Reverse Transcription Kit (Qiagen). qPCR to assess transcript abundance of Irf4, Tbp and Hprt was performed using the following assays: Mm00516431_m1 (Irf4), Mm01277042_m1 (Tbp), and Mm03024075_m1 (Hprt) (Applied Biosystems) using TaqMan Fast Advanced Master Mix (Applied Biosystems) following manufacturer’s instructions. qPCRs were performed using a ViiA 7 Real-Time PCR System (Applied Biosystems).

### Microscopy

Epifluorescence and TIRF imaging were carried out on a Nikon Eclipse Ti microscope with an ORCA-Flash 4.0 V3 digital complementary metal-oxide semiconductor (CMOS) camera (Hamamatsu Photonics) and 100x TIRF objective (Nikon). Illumination was supplied by 405, 488, 552 and 637 nm lasers (Cairn) through an iLas^2^ Targeted Laser Illuminator (Gataca Systems) which produces a 360° spinning beam with adjustable TIRF illumination angle.

Primary B cells from the *in vitro* cultures were stained as described previously before being added to 0.05% Poly-L lysine coated 8-well Lab-Tek imaging chambers (ThermoScientific) and allowed to interact at 37°C. Cells were imaged live. Acquired datasets were analyzed using MATLAB with ImageJ plugin as previously described (Nowosad et al., 2016). Briefly, individual cells were segmented using brightfield and cell surface staining and their fluorescence in the channels showing membrane IgE, IgG1, B220 and CD138 was plotted to allow gating on the desired populations as described for flow cytometry. Mitochondria in gated cells were analyzed by segmenting individual MitoTracker^TM^ Green spots and recording their number and fluorescence.

## RESOURCES TABLE

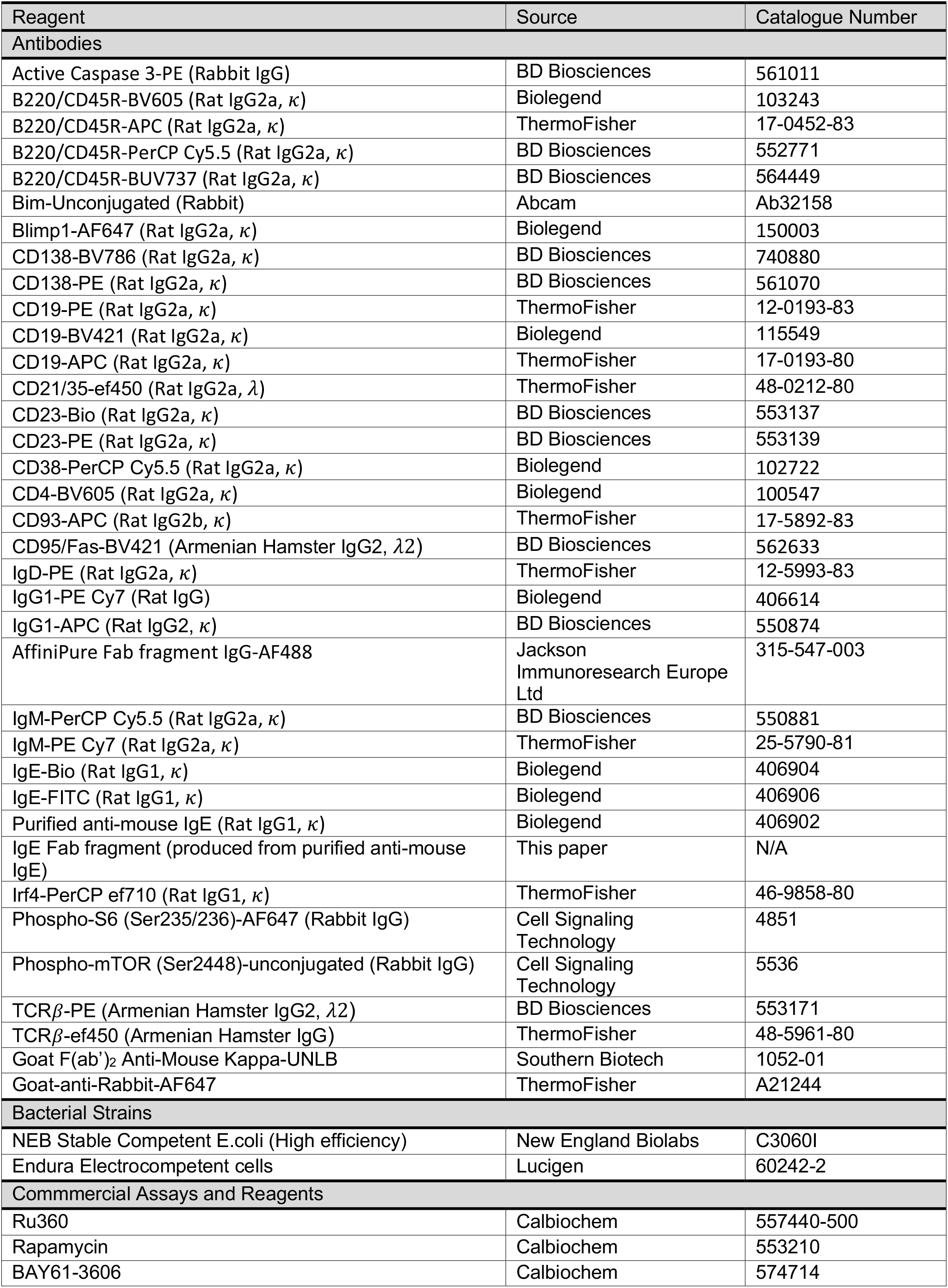
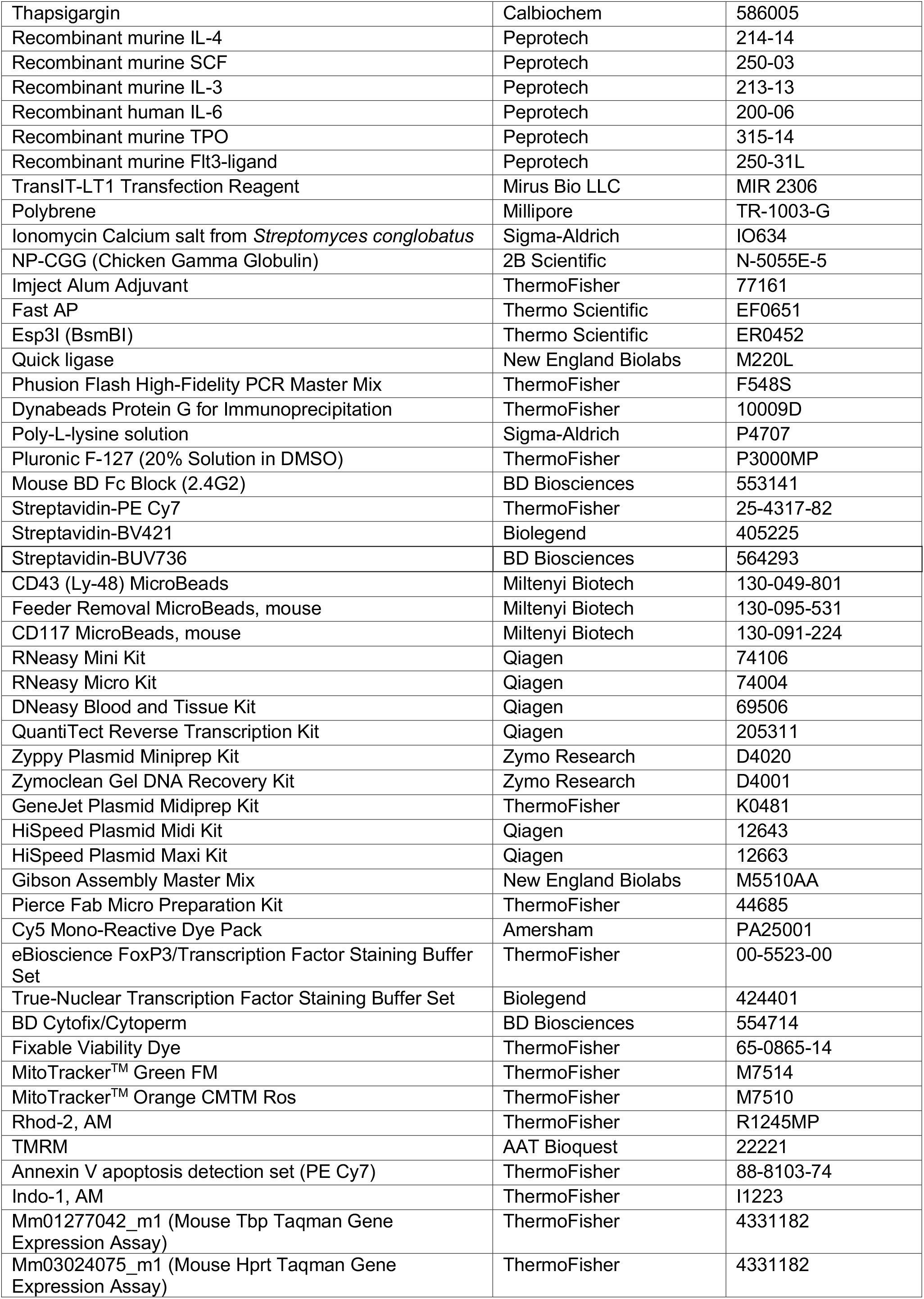
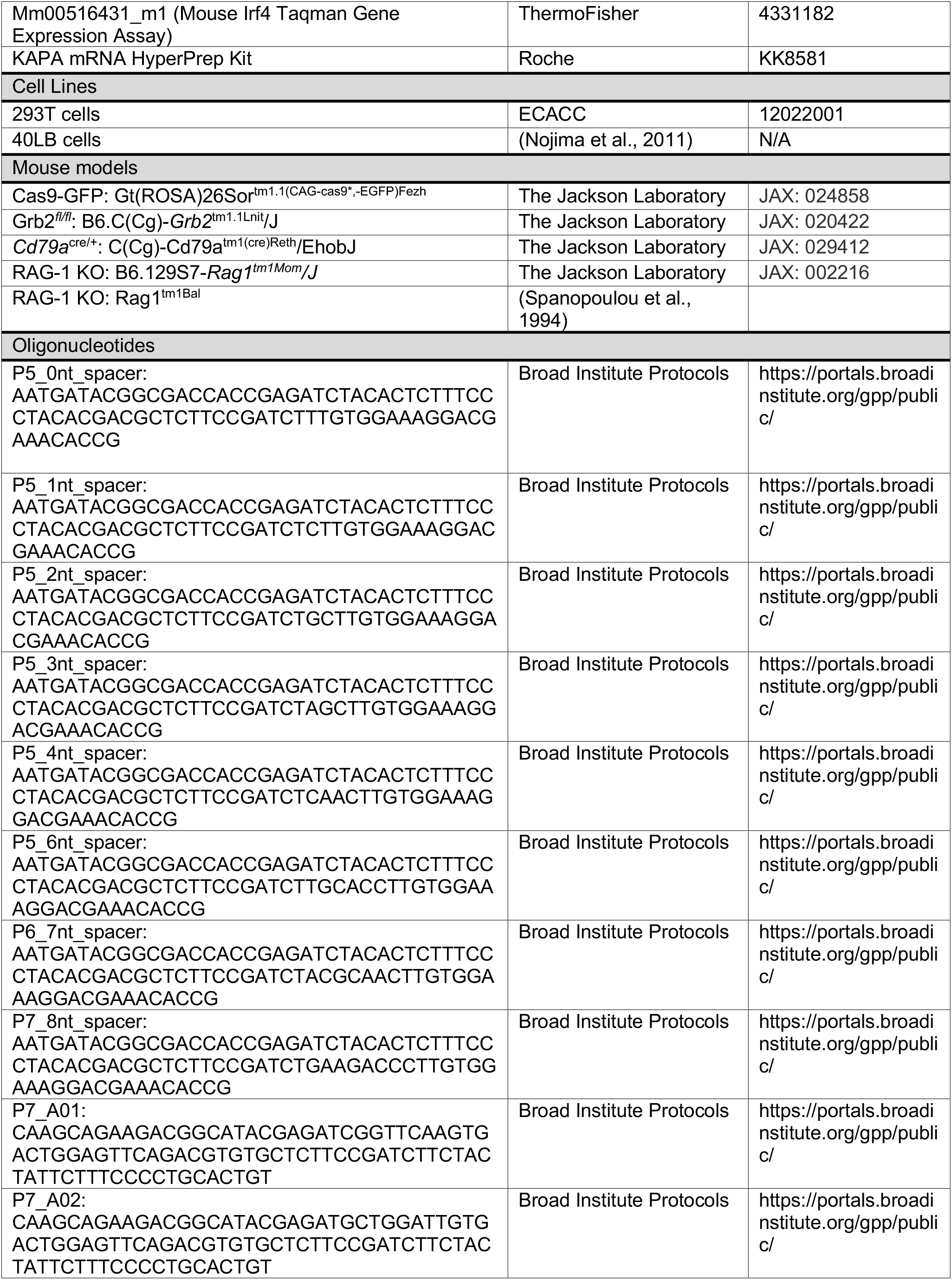
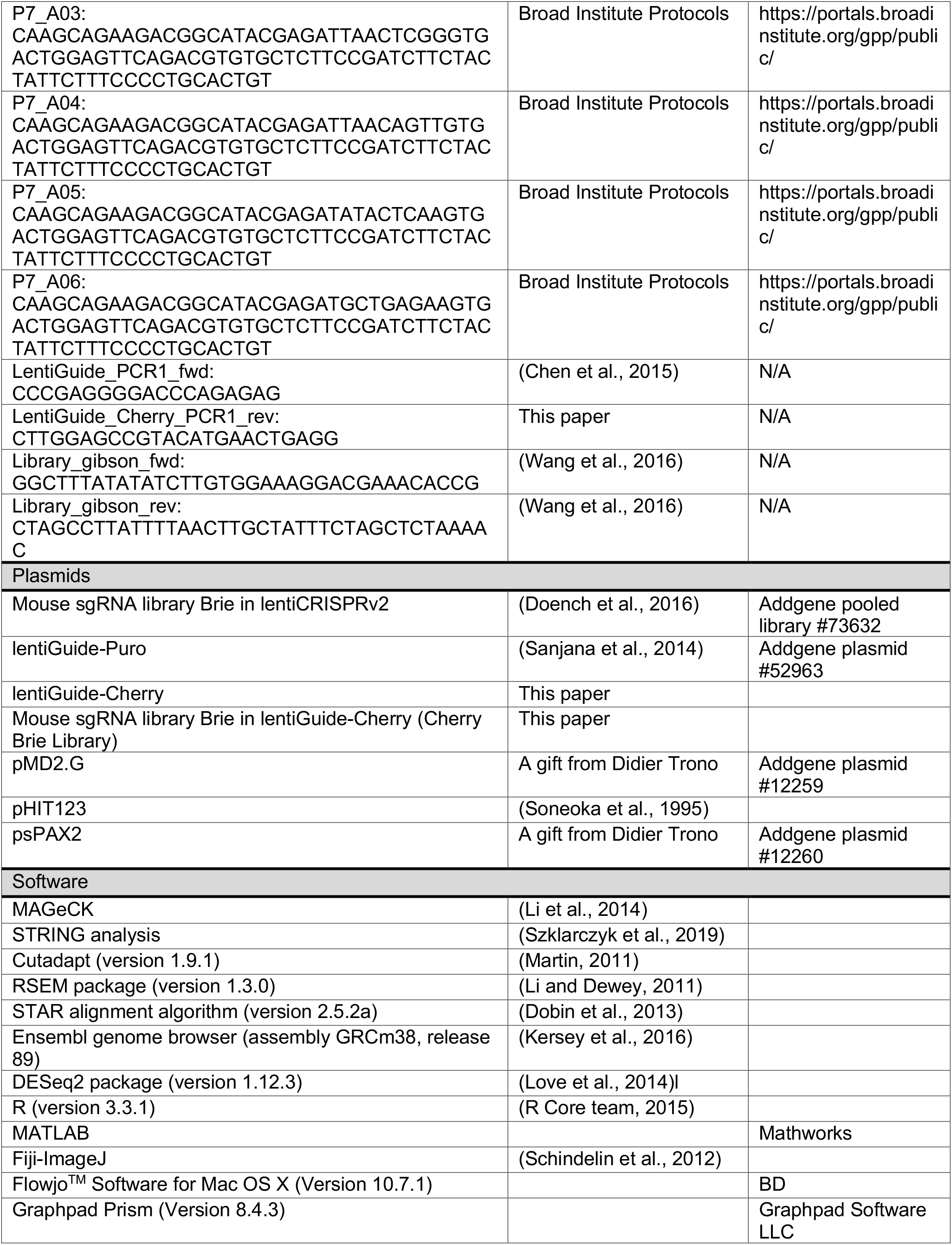

## Notes

### Competing Interest Statement

The authors have declared no competing interest.

